# Delayed suppression normalizes face identity responses in the primate brain

**DOI:** 10.1101/773689

**Authors:** Kenji W. Koyano, Adam P. Jones, David B. T. McMahon, Elena N. Waidmann, Brian E. Russ, David A. Leopold

## Abstract

**Summary:** The primate brain is specialized for social visual perception. Previous work indicates that recognition draws upon an internal comparison between a viewed face and an internally stored average face. Here we demonstrate that this comparison takes the form of a delayed, dynamic suppression of face averageness among single neurons. In three macaque face patches, spiking responses to low-identity morphed faces met with a synchronous attenuation starting approximately 200 ms after onset. Analysis showed that a late-emerging V-shaped identity tuning was sometimes superimposed on linear ramp tuning. This pattern could not be ascribed to repetition suppression within a given session. The results indicate that the brain’s analysis of faces is enhanced through predictive normalization of identity, which increases sensitivity among face-selective neurons to distinctive facial features known to drive recognition.

## Introduction

Humans and other primates rely primarily on vision to extract social information, such as identity and emotional state, from their conspecifics (Leopold and Rhodes, 2010). Primates are highly adapted for face perception, which is reflected in multiple fMRI-defined face patches observed in the inferotemporal cortex of humans, macaques, and marmosets (Hung et al., 2015; Kanwisher et al., 1997; Tsao et al., 2003). The recognition of facial identity is a difficult visual problem that requires extracting the distinctive features of individual faces. Human psychophysical experiments indicate that the brain approaches face recognition by employing an internally stored reference of face averageness, which acts to accentuate feature combinations specific to an individual (Benson and Perrett, 1991; Leopold et al., 2001; Rhodes and Leopold, 2011; Webster et al., 2004). This framework has been formalized as a “face-space” centered on the average face. In this conceptual space, radial trajectories emanating outward from the center are axes of caricaturization that indicate graded levels of individual identity (Valentine, 1991).

While much evidence supports the importance of average face in the perception of face identity, researchers are divided on how this computation may be implemented in the brain (Chang and Tsao, 2017; Freiwald et al., 2009; Kahn and Aguirre, 2012; Latinus et al., 2013; Leopold et al., 2006; Loffler et al., 2005; Panis et al., 2011). A previous electrophysiological study demonstrated that neurons in the macaque inferotemporal cortex showed graded responses to face identity in a manner that resembled the structure of face space, eliciting minimal response to the average face and then systematically increasing their firing rate to more extreme identities (Leopold et al., 2006). Human experiments have shown similar patterns in fMRI responses to faces (Loffler et al., 2005), as well as to human voices (Latinus et al., 2013) and learned shapes (Panis et al., 2011). More recent single unit recordings from macaque face patches also found graded responses to extreme faces, but, unlike the previous studies, did not identify a particular role for average facial features (Chang and Tsao, 2017; Freiwald et al., 2009). The basis of this discrepancy is an open and important question.

An interesting recent perspective on face processing places it in the context of predictive coding and links norm-based coding to hierarchical error signals in the visual system (Issa et al., 2018). In this study, single neurons recorded in the inferior temporal cortex altered their late-phase responses to faces according to the typicality of facial feature configurations. Might the brain’s analysis of facial identity be construed as a type of predictive coding where a stable representation of face averageness serves as the prediction? Mechanistically, this might entail the brain’s active discounting of average facial structural components in order to achieve normalization in the domain of facial identity. Like other examples of normalization in the nervous system, this operation would serve to adjust sensory neurons’ dynamic range to be most sensitive to dimensions that are important for behavior (Carandini and Heeger, 2011). Here we report that single neurons in three fMRI-defined macaque face patches dynamically express such discounting of face averageness through a delayed and synchronous suppression of neural spiking to low-identity face components, resulting in identity normalization.

## Results

We quantitively assessed spiking responses to monkey faces morphed along multiple continuous identity trajectories spanning the center of face space (Fig. 1A). Each trajectory included diminished, full, and exaggerated identity morphs for a given individual (i.e. positive identities), in addition to so-called “anti-faces” (i.e. negative identities). Importantly, anti-faces were generated by extrapolating the morphing transformation beyond the center of face space, such that the features of an anti-face were nominally opposite to those of the original face (Blanz et al., 2000). We recorded spiking responses to 145 photorealistic monkey faces (Fig. 1A-B, Fig. S1) from a total of 296 neurons in three face patches (middle lateral, ML; anterior fundus, AF; anterior medial, AM) of six rhesus macaques (Fig. 1C). We restricted analysis to 186 neurons (103 in AM, 67 in AF and 16 in ML) that showed selectivity for monkey facial identity (*p* < 0.05, two-way ANOVA). Figure 2 shows the response of all the face-selective neurons to the monkey face stimuli. Across the population, neurons exhibited their strongest responses at the extreme identity levels and their weakest responses near the average face. This unprocessed view of all the neurons’ responses to all the morphed monkey faces indicates a special role of the average identity in neural encoding of faces. The next section quantitatively analyzes the nature of the tuning for multiple identity trajectories and in different face patches.

**Figure 1.**
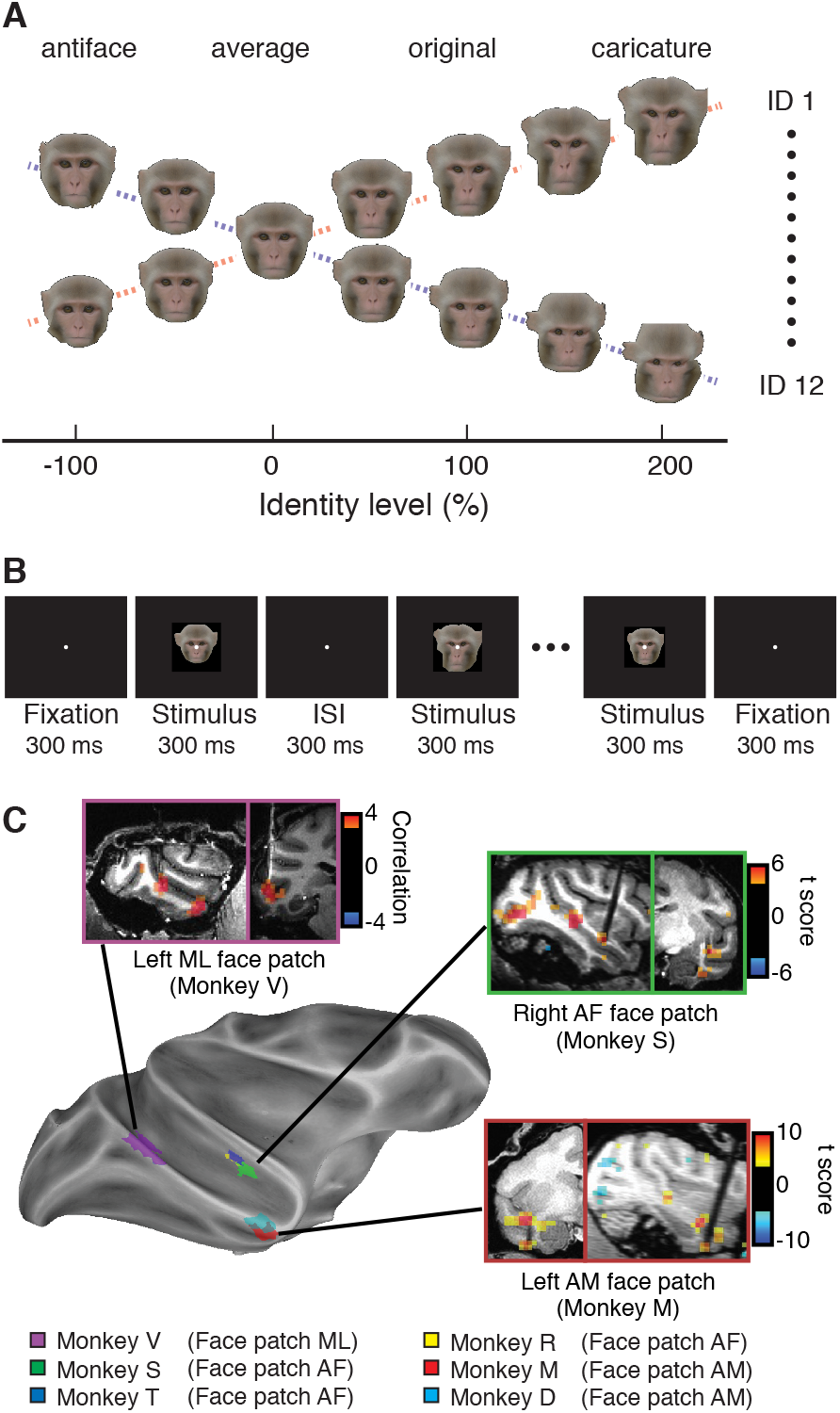
Experimental design. (**A**) Stimulus set. An average face (0% identity level) was first created by averaging 12 monkey faces. Then faces were morphed between the average and each of the 12 original (100%) identities, creating caricatures (200%) and antifaces (−100%) which have exaggerated and opposite facial features of the original, respectively. (**B**) Sequence of passive-viewing task. (**C**) Face patches from where neurons were recorded. See also Figure S1.

**Figure 2.**
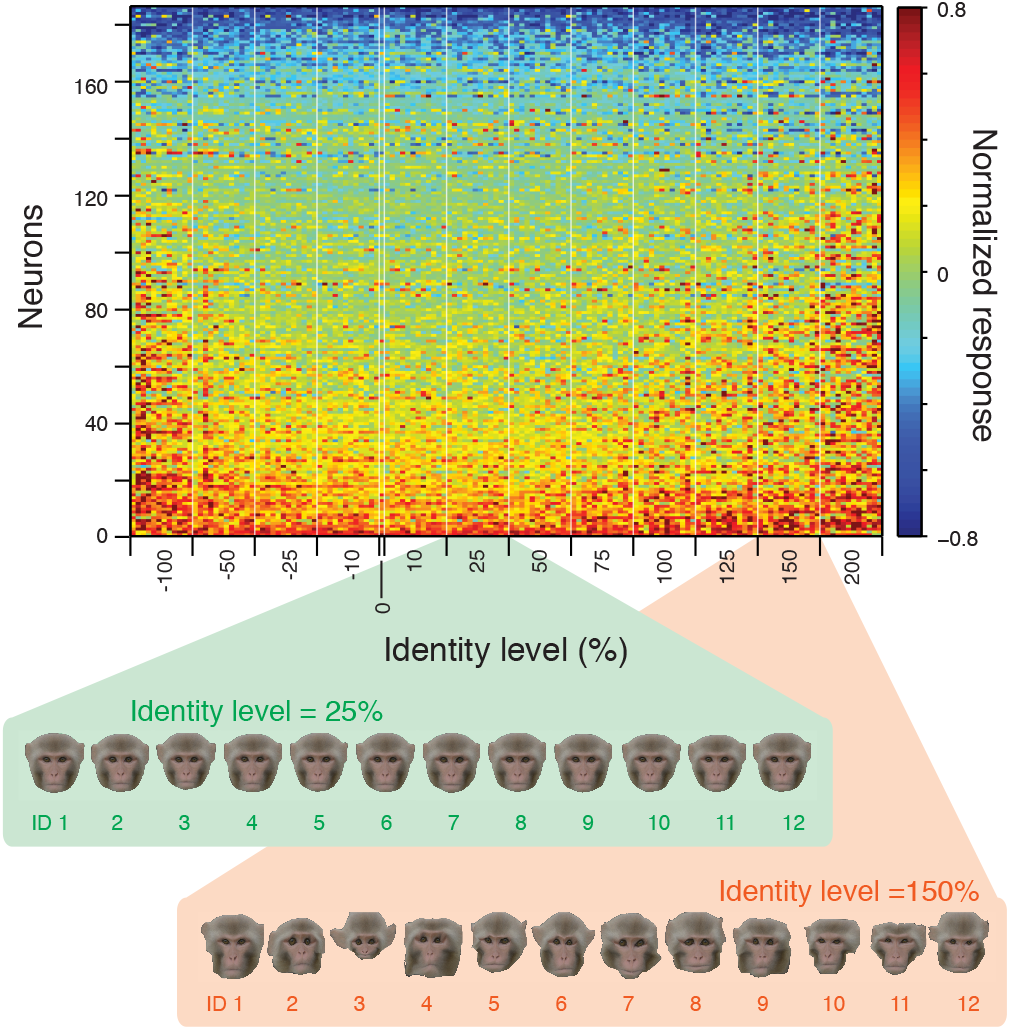
Smallest response to the average face. Normalized responses of all the face-selective neurons (rows, n=186) for all the monkey face stimuli (columns, 145 stimuli). Order of the neurons was sorted by normalized responses that were averaged across the stimuli. Order of stimuli was first sorted by identity trajectory and then sorted by identity level. The high-identity-level (e.g. 150%) faces had more unique facial features specific to each identity trajectory, while low-identity-level (e.g. 25%) faces had less unique facial features. This raw depiction of the data clearly shows that neurons exhibit smaller responses to low-identity-level faces and larger responses to high-identity-level faces.

### V-shaped tuning by face patch neurons

Neurons in each of the three face patches, when tested along single identity trajectories, showed conspicuous dip in response magnitude for the average face. This resulted in a V-shaped identity tuning profile for many neurons (Fig. 3A). A smaller subset of cells showed the opposite profile, with a peak instead of a dip (Fig. S2B). For a given cell, different identity trajectories could exhibit either a slope reversal at the average face (V- or inverse-V-shape), an inflection point (knee-shape), or neither (linear ramp tuning) with different neurons showing varied combinations of identity tuning profiles (Fig S2A-C, S3A-B). For the majority of neurons, averaging across all twelve identity trajectories resulted in clear V-shaped tuning (thick black lines in Fig. 3B and Fig. S2A-C), which was evident across the population of all face-selective neurons (Fig. 3C, Fig. S2D-E). This tuning profile could not arise as the sum or average of linear ramp responses to different identities, but instead depended upon the slope discontinuity at low identity faces that could be extracted by averaging across identity trajectories (Fig. S3F-K). Quantifying these slope changes (Fig. 3D-G), we found that 147 (79.0%) of neurons showed significant slope difference between the caricature and anti-face sides. For each identity, we found a prevalence of V-shaped (21.8 %) and knee-shaped (10.8 %) response patterns (Fig. S3C-D) and 119 (64.0%) of cells had at least one identity trajectory that showed V-shaped or knee-shaped pattern. Further, while individual cells varied in their species selectivity, the same population of cells exhibited similar V-shaped tuning for the human faces relative to the average human face (Fig, S4). As a control experiment, we morphed faces along trajectories between pairs of original identities and confirmed that neurons had no clear V-shaped tuning pattern if the identity trajectory did not pass through the average face (Fig. S5). Together, these observations clearly demonstrate the important role of the average face in the tuning of identity selective neurons, consistent with a previous study (Leopold et al., 2006).

**Figure 3.**
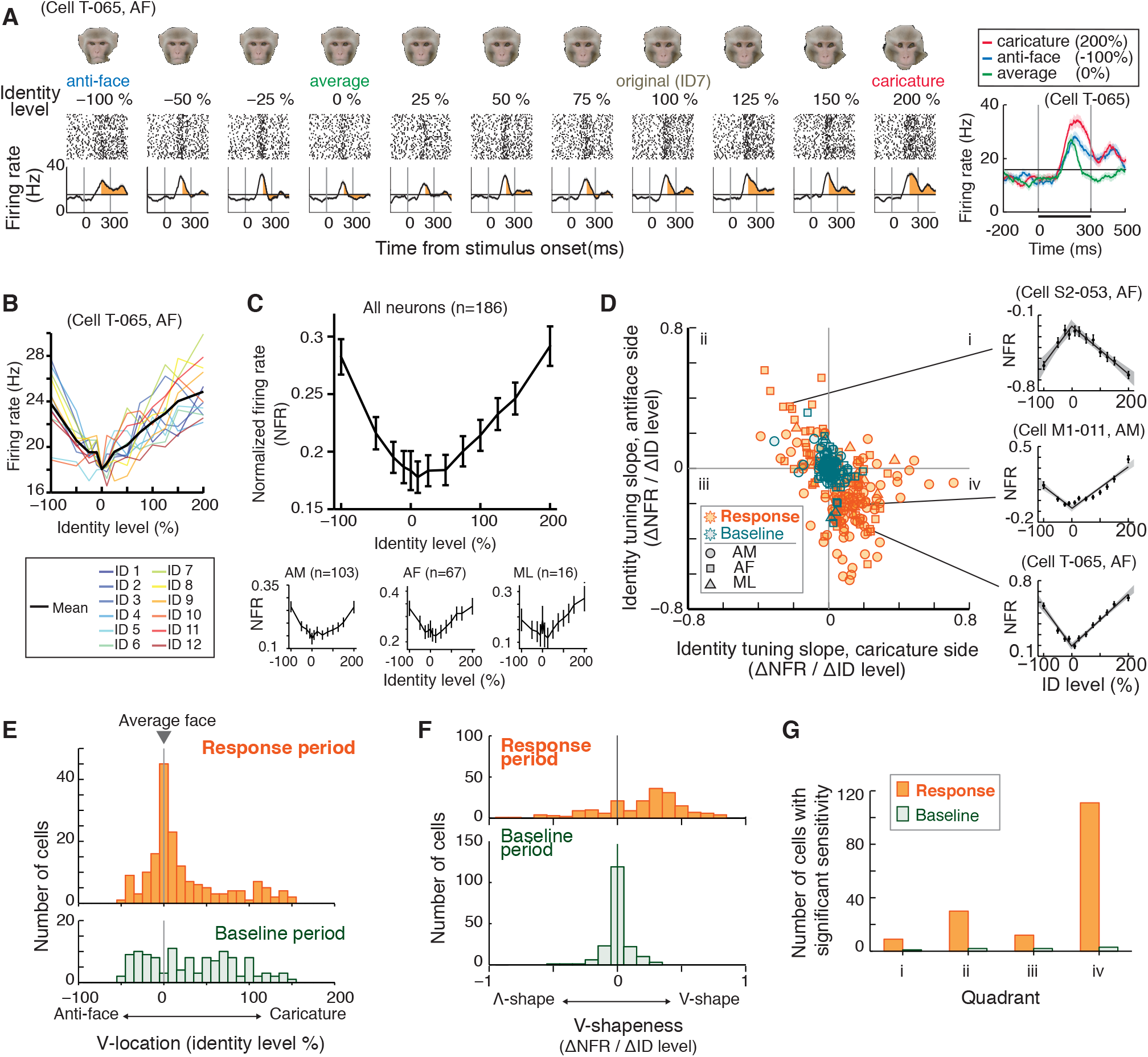
V-shaped tuning by face patch neurons. (**A**) Response of a neuron in an identity trajectory, exhibiting smallest response to the average face. (**B**) Response of the neuron in (**A**) for all the identity trajectories. The mean response showed “V-shaped” tuning pattern along the identity level. (**C**) Mean response of all the face-selective neurons, showing characteristic V-shaped tuning. Insets, mean of each of AM, AF and ML patches. (**D**) Quantification of tuning shape by two linear regression lines. During response period, slopes in caricature side tended to be anti-correlated with slopes in antiface side, indicating V-shaped or Λ– shaped tuning. (**E**) V-location (identity level of crossing point of two regression lines, vertex of the V) distributions during response (*top*) and baseline (*bottom*) period. Many neurons had V-location around 0% identity level (average face) during the response period: 106 (57.0%) neurons showed V-location within ±25% identity level, while only 35 (18.8%) did during the baseline period. (**F**) V-shapeness (difference of regression slope between two lines) distributions during response (*top*) and baseline (*bottom*) period. V-shapeness was significantly biased toward positive values during the response period (*p* < 0.001, Wilcoxon’s sign-rank test). During the response period, 147 (79.0%) neurons showed significantly different regression slopes, while only 5 (2.7%) did during the baseline period. (**G**) Number of neurons with significant sensitivity (significant regression slopes in either side), counted in each quadrant of scatter plot in (**D**). During response period, majority (111, 59.7%) of neurons were in the fourth quadrant (V-shape) and second-majority (30, 16.1%) of neurons were in the second quadrant (Λ-shape). Data are represented as mean ± SEM. See also Figures S2-6.

### Late emergence of V-shaped tuning

We next turned our attention to the temporal evolution of this identity tuning profile. We tracked the time course of the V-shaped tuning using a sliding-window analysis (Fig. 4). Fig. 4A shows the time course of a typical face-selective neuron, showing a large transient response around 100 ms after stimulus onset followed by later sustained response. Note that for this neuron, the V-shaped profile was not clearly evident until late in the response, reaching a plateau only ~200 ms after stimulus presentation. This delayed profile was a shared feature across neurons and clearly evident in the population-averaged response, where V-shaped tuning emerged long after the initial transient response (Fig. 4B). Late tuning for facial identity is consistent with some previous studies reporting higher face identity tuning of neurons in inferotemporal cortex later in the neural response period (Freiwald and Tsao, 2010; Sugase et al., 1999). It is also reminiscent of the more recent results in which the late responses of inferior temporal neurons are modulated by a hierarchical error signal (Issa et al., 2018). Although there were no differences in the initial response latency across identity level, the response magnitude to the low identity faces was abruptly attenuated during the visual response, while that to the high positive and negative identity faces persisted (Fig. 4C, D). Across the population, the number of V-shaped-tuning neurons began to increase around 100 ms and peaked at around 200 ms (55.8 ± 3.5% between 200 ± 50 ms, Fig. 4E), consistent with the shape of the population-averaged tuning. On average, V-shaped tuning emerged 54.8 ± 6.9 ms (mean ± SE) later than firing rate response (Fig. 4F).

**Figure 4.**
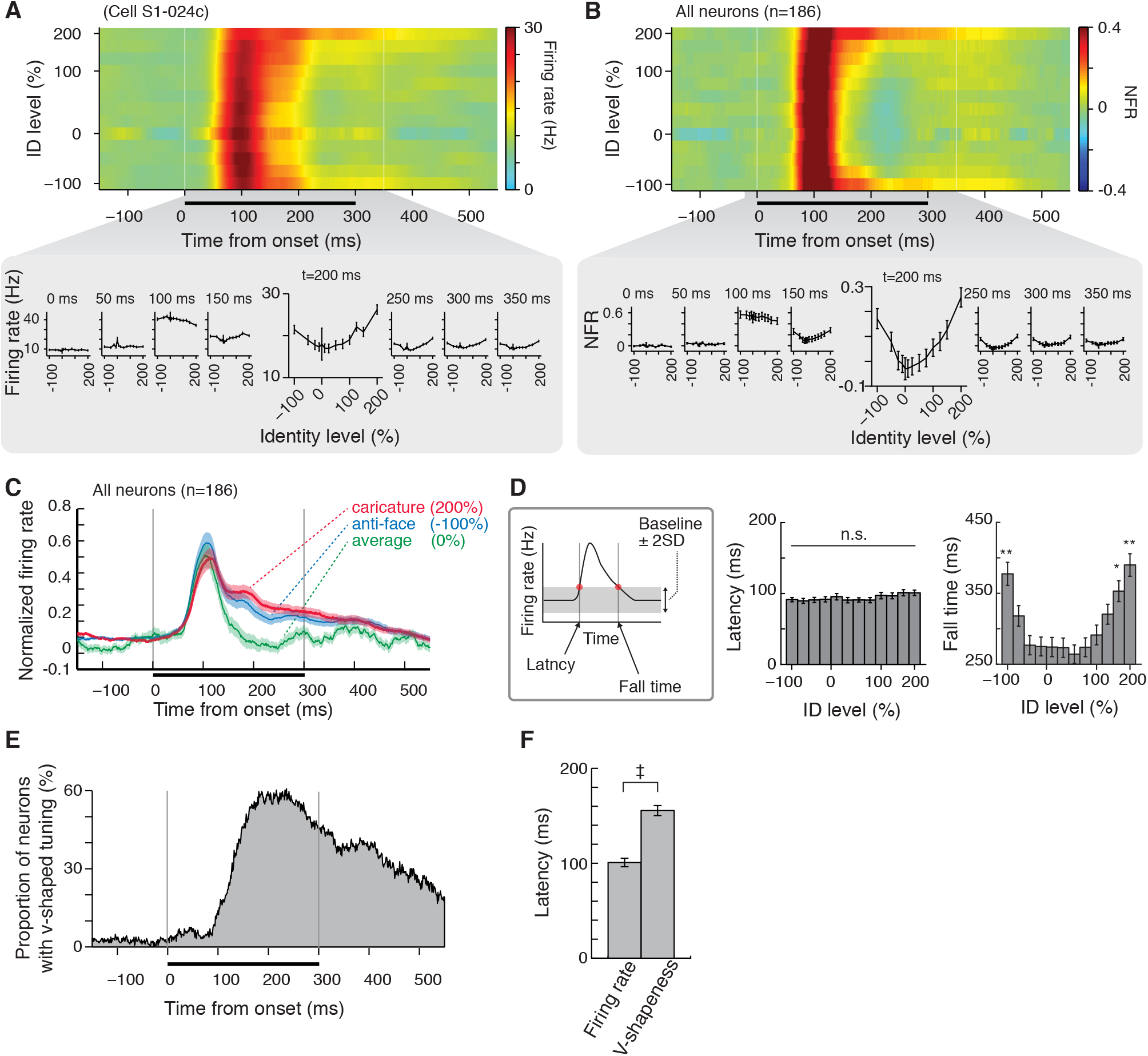
Late emergence of V-shaped tuning. (**A** and **B**) timecourse of the v-shaped tuning pattern in an example neuron (**A**) and population average (**B**). The V-shaped tuning pattern did not clearly observed during the large transient response around 100 ms but slowly emerged during later sustained period that started around 150 ms. (**C**) Mean response timecourses of average, caricatures and antifaces, which diverges in the sustained response period. (**D**) Response latency and fall-time. The response dropped earlier for faces of lower identity levels. *, p<0.05; **, p<0.01 (Tukey’s post-hoc test against faces between −25% to 25% identity level after one-way ANOVA) (**E**) Number of V-shaped tuning cells that increased at later sustained period. (**F**) Latency of V-shapeness, which emerges later than firing rate. ‡, p<0.01 (paired t-test with Bonferroni’s correction). Data are represented as mean ± SEM.

### Neuronal synchrony for the average face

We next asked whether the analysis of spike timing might lend insights into the mechanisms of delayed identity tuning. Our microwire bundles simultaneously recorded local populations of neurons all within a few hundred micrometers (McMahon et al., 2014a). We examined the spiking of neural pairs for their temporal coherence, which is thought to be an index of some aspects of neural coding, including underlying efficient information transmission from a population (Dean et al., 2012; Gregoriou et al., 2009; Koyano et al., 2016). Focusing first on the late visual response, we found that the spike-spike coherence between pairs of neurons (Fig. 5A) was notably stronger for low-identity faces than for more extreme faces (Fig. 5B, Fig. S6A-H), suggesting a larger contribution from the local network during processing of low-identity faces. Coherent spikes were generally coincident in their timing, showing zero-phase coherence (Fig. 5C). Spike-LFP coherence was similarly heightened for low-identity faces (Fig. S6F-H), with spikes most commonly issued in the troughs of the low-γ LFP when oscillatory inhibition should be weakest (Sohal, 2016) (Fig. S6G). Both spike-spike and spike-LFP coherence were highest around the average face and lowest at higher identity levels, resulting in an inverted V-shaped pattern opposite to that of firing rate responses (Fig. 5D). Importantly, this elevated coherence emerged long after the initial visual response and grew coincident with the V-shaped tuning profile (Fig. 5E-F). The late emergence of inverted V-shaped pattern of coherence was also found in neuronal responses for human face stimuli (Fig. S7). These results could not have arisen from a trivial coupling of firing rate and coherence, as we matched the spiking rates in the coherence calculation (see Materials and Methods for details). We interpret the emergence of V-shaped tuning, and its coincidence with heightened spiking synchrony in inverted V-shaped pattern, as a manifestation of broad inhibition during the late response period (Fig. 5G). Increases in spike timing synchrony among excitatory neurons have previously been linked to circuit inhibition (Hasenstaub et al., 2016; Hasenstaub et al., 2005; Sohal, 2016; Sohal et al., 2009; Whittington et al., 2000). In this case, the suppression of the late response was strongest for faces with the lowest identity information. At a systems level, this attenuation of face averageness serves to adjust or normalize the representation of identity to focus on features important for recognition.

**Figure 5.**
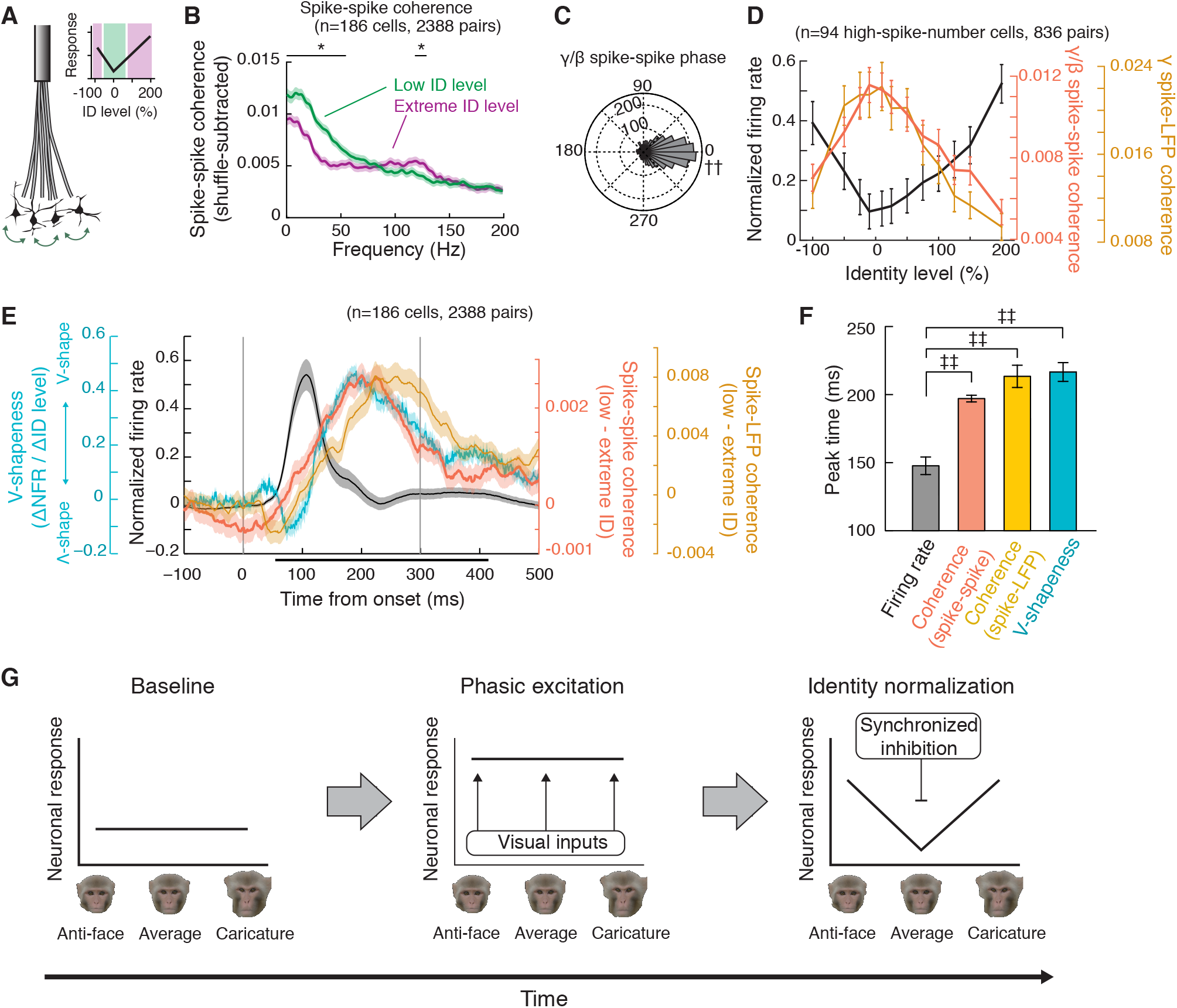
Neuronal synchrony for the average face. (**A**) Simultaneous recording of face patch neurons. Inset, classification of low (green) and extreme (magenta) identity levels. (**B**) Frequency spectrum of spike-spike coherence between simultaneously recorded neuron pairs. The coherence was higher in low identity level than extreme identity level at <50 Hz. *, p<0.05 (paired t-test with Bonferroni’s correction). (**C**) Phase relationships between neuronal pairs, showing coherent firing of spikes. ††, p<0.01 (Rayleigh test). (**D**) Highest coherence for the average face, showing opposite tuning pattern to firing rate. (**E**) Coherence timecourse following v-shapeness timecourse rather than firing rate timecourse. (**F**) Peak time of coherence which is longer than firing rate. ‡‡, p<0.01 (paired t-test with Bonferroni’s correction). (**G**), Model of neuronal mechanism for identity normalization, shaped by general phasic excitation followed by synchronized inhibition for the average face. Data are represented as mean ± SEM. See also Figure S6-7.

### Stable representation of the average face across trials and days

It has previously been suggested that short-term adaptation processes, accrued within a testing session, cause diminished responses to low identify faces (Kahn and Aguirre, 2012). To test whether such short-term adaptation might thus underlie the V-shaped tuning observed in the present study, we examined how this tuning changed within and across sessions (Fig. 6). For this, we capitalized on our microwire bundle electrodes, which routinely maintain the isolation of individual neurons across sessions. For the main population of 186 neurons, each of which was tested across at least two sessions, we found that the tuning shape was consistent on day one and day two, and that tuning corresponding to the first five trials matched that corresponding to the final five trials (Fig. 6A). To investigate this issue over longer timescales, we performed daily, repeated sessions across a population of 13 neurons in the AF face patch that were stably isolated and monitored for 75 sessions spanning three months (Fig. 6B, C). This unique mode of testing gave us the capacity to evaluate whether the neuron’s tuning typically changed over the course of individual sessions, as well as whether the neural tuning changed over the several months of testing period. We did not find evidence for either type of change. Neurons responded weakly to the average face from the very first stimulus presentation of each session (Fig. 6B, C), indicating that such tuning does not arise from within-session adaption (Freiwald et al., 2009; Kahn and Aguirre, 2012). Further, this tuning did not emerge gradually but was present from the monkey’s first viewing of the stimuli and did not change significantly across the 75 sessions. This finding indicates that the V-shaped tuning pattern existed prior to experience with this specific stimulus set, likely because of the subjects’ abundant previous experience with conspecific faces. Lastly, we tested whether the V-shaped tuning might be influenced by the preceding stimulus (Vinken et al., 2018). We found that V-shaped tuning was observed during trials whose preceding stimulus was human faces, which is expected to have minimal adaptation effect onto the tuning shape of monkey faces (Fig. 6D). This again indicated that the V-shaped tuning cannot be explained by within-session adaptation (Freiwald et al., 2009; Kahn and Aguirre, 2012) (Figs. 6A-D). Together, these observations rule out the possibility that within-session adaptation drives the abundant identity tuning around the average face observed in this study.

**Figure 6.**
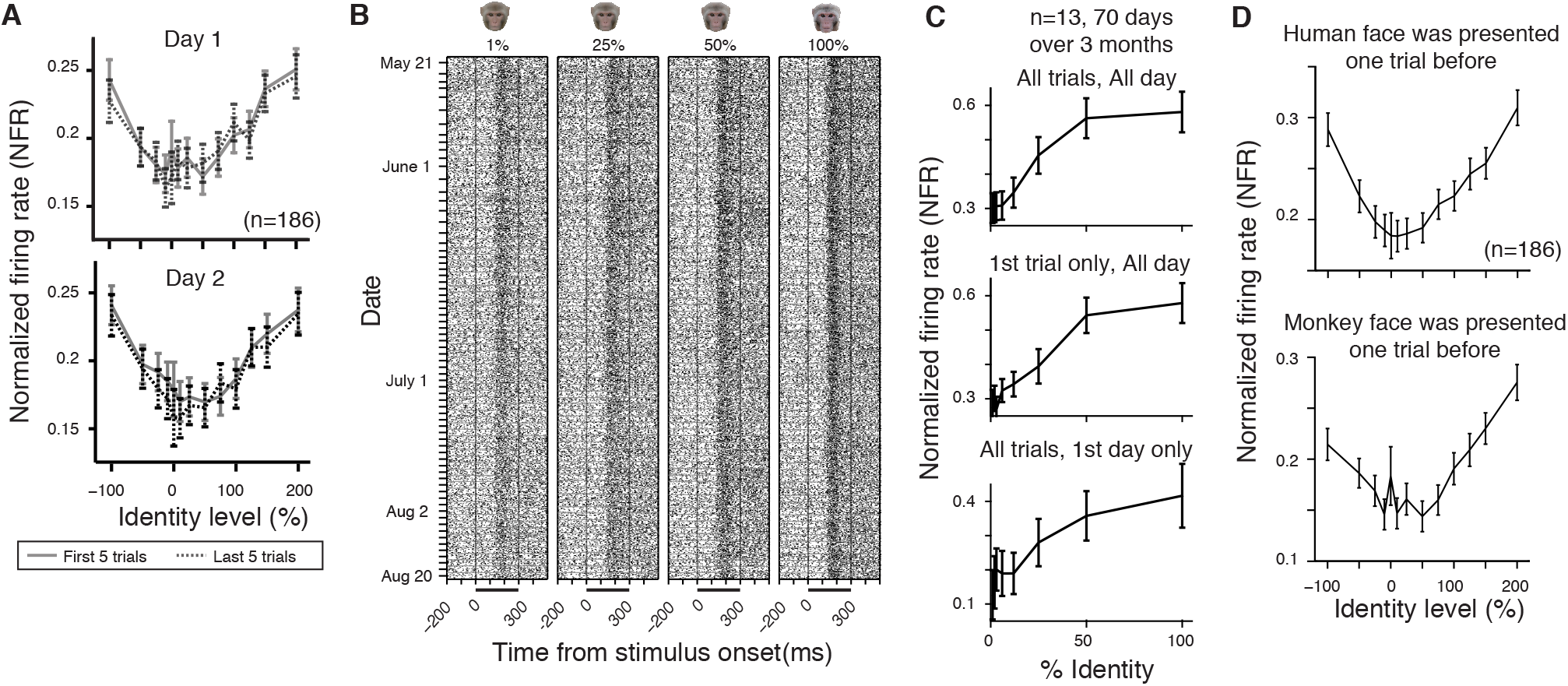
Stable representation of the average face across trials and days. (**A**) V-shaped tuning pattern consistently observed through recording sessions. (**B**) Longitudinal recording from a neuron over 3 months, showing stable tuning pattern along identity levels. (**C**) Tuning pattern consistently observed through the longitudinal recording session. The response was smallest to the average face even in the first trial of each day and first day of the recording, which cannot be explained by adaptation or familiarity effects. (**D**) V-shaped tuning patterns that were observed irrespective of the stimulus type presented one trial before. This again cannot be explained by adaptation. Data are represented as mean ± SEM.

### Identity tuning as a superposition of V-shaped and linear tuning

Previous studies reported linear or ramp tuning for facial features in face-patches (Chang and Tsao, 2017; Freiwald et al., 2009). To estimate contributions of different processes to face identity tuning, we performed regression analysis with a model that combined linear and V-shaped tuning (Fig. 7). Across the population, the face-selective neurons showed a mixed contribution of both components to the tuning shape of each identity trajectory (Fig. 7C). However, those two tuning components exhibited different timecourses (Fig. 7D). The linear tuning developed quickly immediately after stimulus presentation, exceeding V-shaped tuning during 75-201 ms after stimulus onset. In contrast, the V-shaped tuning component gradually increased and became comparable in magnitude to the linear tuning during the sustained response period starting approximately 200 ms after the stimulus onset, consistent with the timecourse of V-shapeness (Fig. 4E and 5E). On average, V-shaped tuning component peaked 35.5 ± 3.7 ms (mean ± SE) later than linear tuning (Fig. 7E). These results indicated that the linear tuning is a major contributor for initial transient response, whereas V-shaped tuning is superimposed upon the linear component during later sustained period, which we postulate to be a normative process serving to enhance the neural sensitivity to identity relative to average face (Fig. 7F).

**Figure 7.**
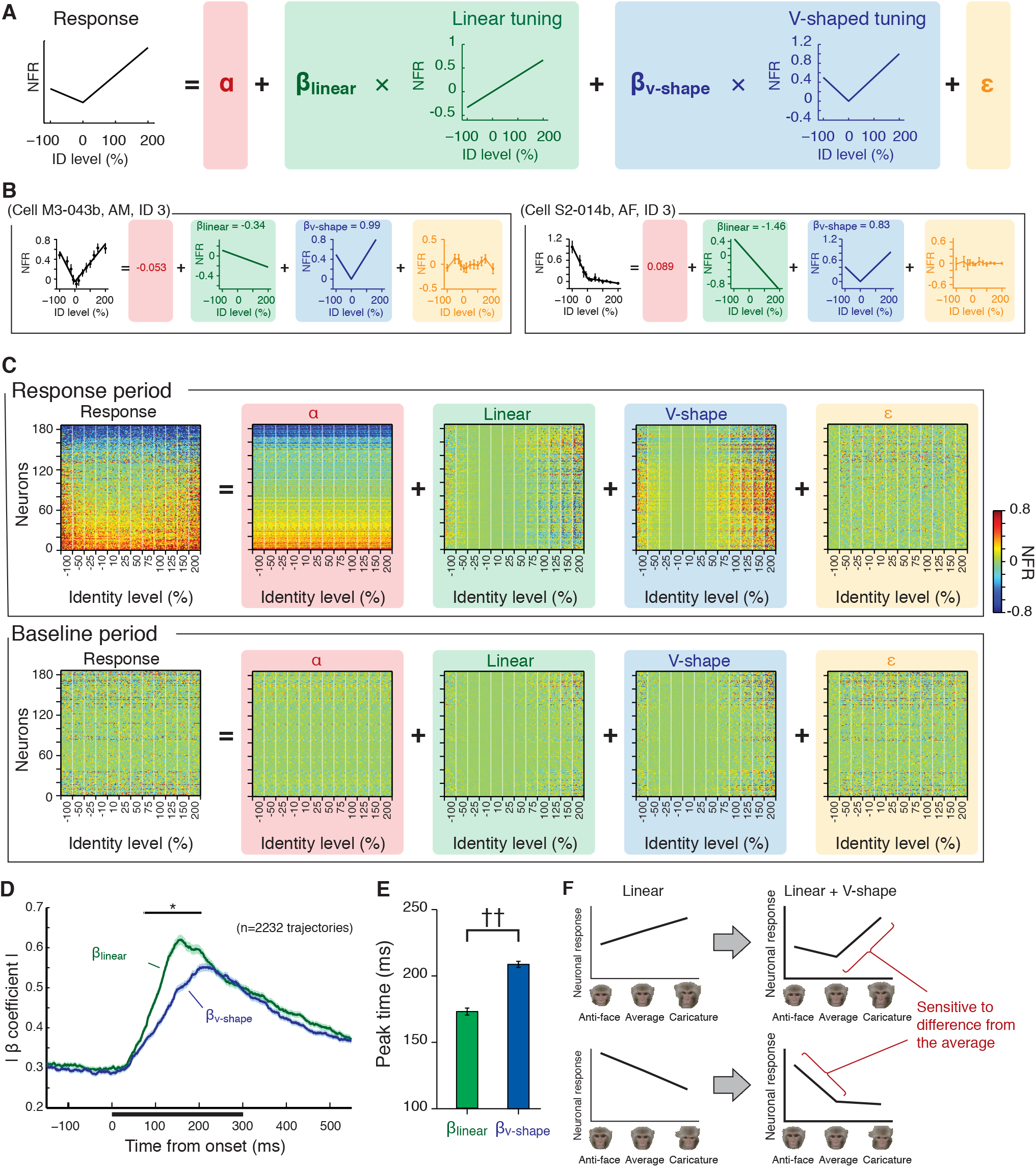
Identity tuning as a superposition of V-shaped and linear tuning. (**A**) Regression model. The neuronal response of each trajectory was modeled with four parameters: intercept *α* (red), coefficient for linear tuning β_linear_ (green), coefficient for v-shaped tuning β_v-shape_ (blue) and error term *ε* (orange). (**B**) Examples of the regression. Neuronal responses were modeled as a summation of linear and v-shaped tunings. (**C**) Regression results for the face-selective neurons (n=186). The responses on the left side was the same one displayed in Figure 2. The responses were represented as a combination of general excitation, linear tuning, V-shaped tuning and error term. (**D**) Time course of the β coefficients. While absolute β coefficient values were comparable during later sustained period after 200 ms, β_linear_ tended to be larger than β_v-shape_ during earlier response during 100-200 ms, indicating larger contribution of linear tuning during the initial transient response. *, p<0.05, paired t-test between the absolute β coefficient values with Bonferroni correction. (**E**) Peak time of the β coefficients showing earlier peak of linear tuning than V-shaped tuning. ††, p<0.001, paired t-test. (**F**) Schematic drawings of how the tuning pattern changed for each identity trajectory. Top and Bottom, examples of different identity trajectories. The tuning was mostly linear at the beginning (*left*), but V-shaped tuning component was added during later sustained period (*right*), leading identity normalization that will make the tuning more sensitive to the difference from the average. Data are represented as mean ± SEM.

## Discussion

We observed the characteristic V-shaped tuning across multiple face patches (Fig. 3C), suggesting that the identity normalization is a common feature in these regions. There was no clear difference in the nature of the V-shaped tuning among face patches, despite a notably high identity selectivity in face patch AM (kurtosis values of 4.81 in AM vs. 3.60 in AF measured across monkey face stimuli, p<0.001, Wilcoxon rank-sum test; see also Freiwald and Tsao, 2010).

Previous studies found a preponderance of linear or ramp tuning, but not tuning around the average face, among face patch neurons for features or principal components signaling identity (Chang and Tsao, 2017; Freiwald et al., 2009). One potential difference from the previous studies is our longer stimulus presentation time (300 ms), which may have given sufficient time for the delayed suppression (Figs. 4, 5) to emerge compared to the short presentations (117-150 ms) in previous studies. Here we found that the linear ramp tuning played a primary role during early transient response, while both the V-shaped tuning and linear ramp tuning played equally important role during later sustained response (Fig. 7D). The initial encoding of facial identity appears to involve a mechanism of rapid linear tuning during the phasic excitation followed by a late-phase, synchronized attenuation of average face information (Fig. 5G, 7F). Consistent with this timecourse, previous studies reported that sensitivity to facial identity was maximized in later sustained response period (Freiwald and Tsao, 2010; Sugase et al., 1999). It is unlikely that the species of the face stimulus is the basis of the difference between the present and previous studies, since we found qualitatively similar effects for both macaque and human faces, and since our previous findings used human faces (Leopold et al., 2006). The method of analysis may also contribute to the observed differences. In the present study, we often summarized individual neuron data by averaging across multiple identity trajectories (Fig. 3, S3). This averaging together of V-shaped trajectories, “knee-shaped” trajectories, and linear ramp-shaped trajectories typically revealed a minimum response at the average face, even though some of the trajectories appeared with no discontinuity there (Fig. S3).

### Identity normalization of faces

Normalization is an operation found in multiple areas of the brain including early visual cortices (Carandini and Heeger, 2011), where range of neuronal responses is dynamically adjusted through the normalization process to maximize neural sensitivity. Here, we found a similar normalizing operation of face identity in higher visual areas of the inferotemporal cortex, which maximized neural sensitivity for the difference from the average face (Fig. 7F). Consistent with the psychological concept of norm-based face coding (Benson and Perrett, 1991; Leopold et al., 2001; Rhodes and Leopold, 2011; Valentine, 1991; Webster et al., 2004), the identity normalization can increase neuronal sensitivity to faces in a familiar group of people (Valentine, 1991). While early visual cortices utilize visual input itself for the normalization of the visual input (Carandini and Heeger, 2011), the identity normalization relies on subjects’ past experiences which works as a top-down modulation signal, similarly to normalization of attention through a top-down signal (Reynolds and Heeger, 2009).

### Acquisition of the norm

How the brain gradually gathers information about important stimuli such as faces to generate norms for certain visual problems is currently a matter of conjecture. For faces, one possibility is that this information is learned and stored external to the visual system proper, perhaps in an area such as the perirhinal cortex that shares dense reciprocal connections with the inferior temporal cortex (Lavenex et al., 2002). Recent work showing that a face-specific subregion of the perirhinal cortex responds selectively to familiar faces (Landi and Freiwald, 2017) suggests that this area might provide an external feedback signal to face patches, perhaps normalizing responses according to face averageness. Another possibility is that the disparate face patches exhibit a parallel learning of facial structure, mainly through experience during development, which they then apply locally through local inhibitory mechanisms. Future work may address whether this mode of information encoding is unique to faces, or whether it may also pertain to other stimuli such as color, shape, or bodily postures (Bell et al., 2009; Lafer-Sousa and Conway, 2013; Popivanov et al., 2012; Srihasam et al., 2012). Likewise, an intriguing possibility for future study is that this mechanism for face coding in primates reflects a more general principle of how brains apply gradually learned statistics of important stimulus categories, and particularly those related to the perception of conspecifics, an aspect of visual cognition thought to be critical for sexual selection (Bradley and Mundy, 2008; Leopold and Rhodes, 2010; Rhodes, 2006).

## Materials and Methods

### Subjects

Six rhesus monkeys (*macaca mulatta*, four females and two males, weighing 5.5 - 11.3 kg) were used in this study. All animals were surgically implanted with an MRI-compatible head post, and with chronic microwire electrode bundles in a face patch (Fig. 1C; AM face patch for two animals, AF face patch for three animals and ML face patch for one animal) which was functionally localized using a standard fMRI block design (McMahon et al., 2014b; Russ and Leopold, 2015) and/or naturalistic movie watching paradigm (Russ and Leopold, 2015). The apparatus and surgical implantation protocol have been described in detail previously (McMahon et al., 2014a). All surgeries were performed under aseptic conditions and general anesthesia under isoflurane, and animals were given postsurgical analgesics and prophylactic antibiotics. During participation in the recording experiment, the animals were on water control and received their daily fluid intake during their testing (see below). Each subject’s weight and hydration level were monitored closely and maintained throughout the experimental testing phases. All the experimental procedures and animal welfare were in full compliance with the Guidelines for the Care and Use of Laboratory Animals by U.S. National Institutes of Health and approved by the Animal Care and Use Committee of the U.S. National Institutes of Mental Health / National Institute of Health.

### Behavioral task and visual stimuli

The monkeys sat in a primate chair in front of an LCD/OLED monitor with their head position stabilized by means of an implanted head post. They were required to maintain their gaze on a fixation point of 0.2° × 0.2° at the center of the monitor through a trial. In each trial, visual stimuli of 7° × 7° monkey face or 4° × 6.4° human face were presented for 300 ms in pseudo-random order followed by a 300 ms inter-stimulus interval (Fig. 1B). The monkeys were rewarded with fruit juice for successfully maintaining fixation within a window of 1.5° - 2°, while their eye position was monitored using an infrared video-tracking system (EyeLink II; SR Research). Stimulus presentation, eye position monitoring, and reward delivery were controlled by MonkeyLogic software (Asaad et al., 2013), NIMH MonkeyLogic Software (Hwang et al., 2019) or custom software courtesy of David Sheinberg (Brown University, Providence, RI) running on a QNX computer in combination with another machine that run the psychtoolbox (Kleiner et al., 2007) in Matlab (Mathworks). The monitor (either a ViewSonic 18” LCD monitor or LG 55” OLED monitor) was placed 91 or 90 cm in front of the monkey. Timing of stimulus presentation was recorded by a photodiode sensor that received signal from a small white square displayed on a corner of the screen at the same time of stimulus presentation.

The photo-realistic face stimuli were generated by morphing between face images (Fig. 1A, Figs. S4 and S5) using a face-morphing software (Fantamorph, Abrosoft). Monkey faces were created based on twelve monkey face photographs of the NIH colony, prepared and provided by Dr. Olga Dal Monte, and human faces were created based on twenty four human faces selected from the FEI face database (https://fei.edu.br/~cet/facedatabase.html). We first created the average face that resides at the center of the face space and approximates a norm (Valentine, 1991), by a morphing algorithm using point by point correspondence of the individual face. Then the stimuli were created by morphing along face-space trajectories between the average and original faces, that is to say an identity trajectory (Blanz et al., 2000). By extending the identity trajectories past the original faces, caricature faces that had exaggerated features were created. We also extended the identity trajectories away from the average, creating so-called anti-faces that had “opposite” features of the original faces. The morphing percentage was taken as the “identity level”, in which 0% identity level corresponded to the average face and 100% identity level corresponded to the original full identity faces. We assigned negative identity levels of <0% to anti-faces. Stimuli were generated at identity levels of −100, −50, −25, −10, 0, 10, 25, 50, 75, 100, 125, 150 and 200%. In total, 145 monkey faces and 289 human faces were created and used. In this study, we morphed the faces based on shape information and applied average texture to all the images to exclude confounding effects of the texture smoothness which could be maximized at the norm stimulus. For the experiment in Figure 6, we generated stimuli along 8 identity trajectories at the identity levels of 1, 2, 5, 10, 25, 50, 75 and 100%, based on both shape and texture information. For the experiment in Figure S5, we morphed between pairs of original identities at the identity levels of 0, 10, 25, 50, 75, 90 and 100% (for this morphing, we defined 0 and 100% identity levels as one of paired original faces). In addition to the face stimuli, 12 object and pattern images were also shown to ensure that the same neurons were held across experimental sessions on different days.

### Electrophysiology

Extracellular neuronal signals were recorded with 32, 64 or 128 chronically implanted NiCr wires that permitted tracking of individual neurons over multiple recording sessions (McMahon et al., 2014a; McMahon et al., 2014b). The microwire electrodes were designed and initially constructed by Dr. Igor Bondar (Institute of Higher Nervous Activity and Neurophysiology, Moscow, Russia) and subsequently manufactured commercially (Microprobes). The recorded neuronal signals were amplified and digitized at 24.4 kHz in a radio frequency-shielded room by PZ5 NeuroDigitizer (Tuker-Davis Technologies), and then stored to an RS4 Data Streamer controlled by an RZ2 BioAmp Processor (Tucker-Davis Technologies). A gold wire inserted into a skull screw was used for ground. Broadband signals (2.5-8 kHz) were collected, from which individual spikes were extracted offline using the WaveClus software (Quiroga et al., 2004) after filtering between 300 and 5000 Hz. Local field potentials (LFPs) were extracted off-line from the same broadband signals by zero-phase, bi-directional fourth-order Butterworth band-pass filtering between 3-200 Hz. Event codes, eye positions and a photodiode signal were also stored to a hard disk using OpenEX software (Tucker Davis Technologies).

The method for longitudinal identification of neurons across days was described in detail previously (Bondar et al., 2009; McMahon et al., 2014a; McMahon et al., 2014b). Briefly, spikes recorded from the same channel on different days routinely had closely matching waveforms and interspike interval histograms, and were provisionally inferred to arise from the same neurons across days. This initial classification based purely on waveform features and spike statistics was tested against the pattern of stimulus selectivity and temporal structure of the neurons’ firing evoked by visual stimulation. Guided by our previous observations that neurons in inferotemporal cortex respond consistently to statically presented visual stimuli across days and even months (Bondar et al., 2009, McMahon et al., 2014), we used the distinctive visual response pattern generated by isolated spikes as a neural “fingerprint” to further disambiguate the identity of single units over time. The longitudinal aspect of the recording allowed us to collect neuronal responses for a large number of stimuli. By spending 2.96 ± 0.89 days, we recorded neuronal responses 19.91 ± 5.65 times for each of 536 stimuli, resulting in 10673.3 ± 3026.2 trials in total. In addition, we recorded responses from single neurons for 75 days, which enabled us to evaluate within-session adaptation for the stimuli (Fig. 2E-G).

### Data analysis

Stored neuronal response data were analyzed offline with MATLAB software (Mathworks, MA). All the data in the text was expressed as mean ± S.D. unless otherwise stated. Error bars in figures are standard error unless otherwise stated. To evaluate V-shaped tuning, firing rate responses of neurons to each stimulus were calculated from 200 to 500 ms after the stimulus onset. Stimulus selectivity of the responses were evaluated for each neuron by two-way analysis of variance (ANOVA) [two factors: identity trajectories (identity of 12 original monkey faces or 24 original human faces of the trajectory) and identity level (12 identity levels ranging from −100% to 200%, excluding 0% identity face that is shared across all the trajectories)]. Neurons which showed one or more main effect and/or interaction between factors (p < 0.05 after Bonferroni correction) were considered as face-selective and further analyzed.

Mean tuning for identity level were calculated by averaging across all the identity trajectories for each neuron (Fig. 3B). Population-averaged tuning was calculated by averaging normalized firing rate response (NFR), which was calculated from the maximum and baseline responses of each neuron (Fig. 3C). Baseline response was calculated from 200 ms before the stimulus onset to 50 ms after the stimulus onset across all the trials of each neuron. Maximum response was defined as the response to the stimulus that elicited largest response, either excitation or suppression, as compared to the baseline response. The response for each stimulus was normalized by subtracting the baseline response and then dividing it by the absolute difference between the baseline and maximum response of the neuron, resulting in normalized firing rate value that ranges from −1 to 1 (−1 or 1 corresponded to the maximum response and 0 corresponded to the baseline response). Neurons whose mean response was less than the baseline were considered as suppressive neurons, and the sign of their normalized firing rates was inverted before calculating population-averaged response, since the tuning shape tended to be inverted for those neurons (R = 0.31 between normalized mean response and V-shapeness).

For each neuron, the V-shaped tuning pattern was quantified by modeling the tuning with two regression lines, which included four parameters: anti-face side (left side) slope, caricature-face side (right side) slope, identity level of the vertex (V-location, x-axis value of a crossing point of two regression lines) and NFR of the vertex (y-axis value of the crossing point). The optimal modeling parameters were estimated by least-squares technique with maximum of 2,000 iterations. If parameters were not estimated after 2,000 iterations, or if the estimated vertex was located too right or left where not enough (fewer than two) identity levels were available to estimate either the right or left side slope, the initial value was changed and the estimation process was repeated again. If the parameters were not successfully estimated after a total of 100,000 iterations, we dismissed the two-line model and adopted a single-line linear regression for that neuron. Of 186 face-selective neurons, response activities of 172 (92.5%) neurons were modeled with the two-line model, while baseline activities of 121 (65.1%) neurons were modeled with the two-line model. The estimated tuning slopes were represented as a change of NFR per 100% identity level. The tuning pattern of each identity trajectory was quantified by modeling with two regression lines using three parameters: anti-face side (left side) slope, caricature-face side (right side) slope and NFR of the vertex. Identity level of the vertex in each trajectory was fixed at 0% to prevent overfitting due to fewer data points. The optimal modeling parameters were estimated by least-squares technique as modeling for each neuron. The average face, whose identity level was 0%, was not included in the regression to avoid duplicated contribution to multiple identity trajectories since it was shared across different identity trajectories. In Figure 7, each identity trajectory was also modeled by combination of linear ramp tuning and V-shaped tuning (see Fig. 7A for the model function). The basis functions of the linear and V-shaped tuning were both designed to cross the origin [0, 0] and to cover a range of ±0.5 of normalized firing rate in the y-axis. V-shapeness was defined as the difference of slopes between caricature and antiface sides.

Spike trains were smoothed by convolution with a Gaussian kernel (σ = 30 ms) to obtain spike density functions (SDF) for each stimulus (Fig.3A). To evaluate the temporal dynamics of firing rate response and regression slopes (Fig. 4), firing rates were calculated with a sliding window of ±25-ms width moving in 1-ms steps. To evaluate the temporal dynamics of linear and V-shaped tuning for each identity trajectory, sliding window of ±50-ms width was used (Fig. 7D, E). Firing rate responses at each time point were normalized in a similar manner to the normalized firing rate as described above, with the baseline response which was calculated between 200 ms before and 50 ms after the stimulus onset and the maximum response which was calculated between 200 and 500 ms after the stimulus onset. The onset latency was defined as the time after the stimulus onset when the firing rate exceeded ±2 S.D. of the firing rate during the 250-ms baseline period for 10 consecutive bins and reached to 3 S.D. of the baseline firing rate at least once. The fall time was defined as the time from the stimulus onset when the firing rate decreased below ±2 S.D. of the firing rate during the baseline period for 10 consecutive bins and reached to 1 S.D. of the baseline firing rate at least once. The regression slopes were estimated at each time point of the normalized firing rate timecourse in the same manner of the two-line modeling described above. To evaluate dynamics across firing rate response, neuronal tuning and coherence (Fig. 5E), we calculated peak time since it depends less on time window width, which needed to be relatively large to ensure a high enough signal-to-noise ratio for coherence computation. Peak time was also used for comparing β coefficients between linear and V-shaped tuning (Fig. 7E) which needed relatively large time window for regression of each identity trajectory from fewer number of trials. The peak time was defined as the time point after the stimulus onset when the timecourse reached its maximum value between 0 to 400 ms after the stimulus onset.

We calculated spike-spike and spike-LFP coherence to evaluate synchronized computations that may be processed across neural circuits. Coherence spectrum was estimated with multi-taper spectral methods (Mitra and Bokil, 2008; Mitra and Pesaran, 1999) using Chronux toolbox (http://chronux.org) and CircStat toolbox (Berens, 2009). For comparisons between low and extreme identity faces, five Slepian taper functions were used with a ±75 ms time window with ±10 Hz resolution. Faces whose absolute identity level was equal to or less than 50% were classified as low identity faces, and faces whose absolute identity level was equal to or larger than 75% were classified as extreme identity faces. Eleven Slepian taper functions were used with a slightly larger window, ±100 ms and ±15 Hz resolution, for comparison across identity levels which had fewer trials per data point. In addition, we excluded neurons whose mean spike number per identity level was less than 200 (for human faces in Fig. S7 and for non-spike-remove control analysis in Fig. S6J, we excluded neurons whose spikes per identity level was less than 400) for the comparison across identity levels. Coherence was calculated during the later sustained response phase, starting at 150 ms after stimulus onset. Coherogram and coherence timecourse were computed with sliding window of ±75-ms width moved at 1 ms steps. Spike-spike coherence was calculated between all the possible neuronal pairs that were simultaneously recorded in the same session. For calculation of spike-LFP coherence, LFP signals were averaged across all the recording channels of the microwire bundles except for noisy or dead channels. To avoid potential contamination of LFP signals by spike waveform, the channel that recorded the spikes for a given coherence calculation was also excluded from the averaging of LFPs. Based on coherograms (Fig. S6H) and frequency spectrums of coherence (Fig. 5B), we averaged coherence and phase in low-γ/β range (15-50Hz, for spike-spike coherence) and low-γ (30-50 Hz, for spike-LFP coherence) for comparisons across identity levels and between low and extreme identities. The number of trials was balanced across conditions: the difference in number of trials was 1.52 ± 0.7% for comparisons between low and extreme identity levels and 2.9 ± 2.2% for comparisons across identity levels. Thus potential effect of bias from different sample sizes was minimized.

We examined the effect of identity levels on the coherence, which also affected firing rate responses of each neuron. Although coherence is designed to be invariant over differences of firing rate by normalization (Mitra and Bokil, 2008; Mitra and Pesaran, 1999), some aspects of the coherence measure, like signal-to-noise ratio, could be influenced by firing rate. To rule out any potential contribution of firing rate differences to coherence, we matched firing rate across conditions before coherence calculation by randomly dropping spikes (Dean et al., 2012; Gregoriou et al., 2009; Mitchell et al., 2009). During the comparisons between low and extreme identities, for each neuron, we randomly dropped spikes of one condition that showed higher firing rate until it matched with the other condition. During the comparisons across identity levels, firing rates were matched to an identity level that showed minimum firing rate response. For calculating coherograms and coherence timecourses, we separated timecourse into 200-ms-width bins and matched firing rates in each bin. Matching of firing rate did not affect essential aspects of coherence of our data; even without any firing rate matching, low identity faces elicit higher γ/β coherence during late phase of the response (Fig. S6I-L). Matching of firing rate reduced absolute coherence values, especially at high-γ range over 100Hz in higher identity level conditions which elicit larger firing response (Fig. 5B compared with Fig. S6I), suggesting that the observed coherence at high-γ range might be partially reflecting firing rate response.

To control coherence arising from stimulus-locked response, trial-shuffled coherence was also calculated. For each pair of neurons and each neuron-LFP pair, the order of trials was randomly shuffled within each stimulus in a neuron while maintaining the trial order of the other signal (the other neuron or LFP), and then coherence was computed based on the trial-shuffled spike trains. The trial-shuffled coherence was created 1000 times, and their mean was subtracted from the original coherence to produce trial-shuffle-subtracted coherence. For controlling coherogram and coherence time course data, one-trial shifted control was subtracted from the original to produce a trial-shift-subtracted coherogram and coherence timecourse. The trial-shuffled or trial-shifted control were subtracted in all the coherence/coherogram data displayed in Figures, except for the “Raw” coherence in Figs S6B,F,J and Fig S7D.

### Long duration recordings to investigate role of visual adaptation

We exploited the stable recordings of our microwire electrodes to track a population of 13 neurons in the AF face patch across 75 sessions spanning three months beginning with the first presentation of each stimulus. As above, we used the selectivity “fingerprint” to establish the identity of individual neurons for tracking during this period, a method that has been used effectively previously (McMahon et al., 2014b). For this data collection, only positive face identities, between 0 and 100% identity were used, with no anti-faces or caricatures. Collecting data with the same stimulus set across days, from the first presentation of each stimulus, allowed for the evaluation of changes both within and between sessions. The 75 sessions allowed us to draw selectively from the first presentation of each stimulus during the day, averaging this first presentation across sessions, to determine whether neural responses to the first presentation differed critically from that those to later presentations. This allowed us to test the hypothesis that the V-shaped tuning of face-selective neurons might arise due to short-term visual adaptation.

## Acknowledgments

We would like to thank David Yu, Charles Zhu and Frank Ye for assistance with fMRI scanning, Julie Hong, Katy Smith, George Dold and David Ide for technical assistance and development, Olga Dal Monte and Carlos E. Thomaz for providing face stimli, Vasileva Liubov, Igor Bondar, David Godlove, Bruno Averbeck, Teppei Matsui and Toshiyuki Hirabayashi for helpful discussions. Functional and anatomical MRI scanning was carried out in the Neurophysiology Imaging Facility Core (NIMH, NINDS, NEI). This work utilized the computational resources of the NIH HPC Biowulf cluster http://hpc.nih.gov). This work was supported by the Intramural Research Program of the National Institute of Mental Health (ZIAMH002838 and ZIAMH002898) to D.A.L. K.W.K. was supported by the fellowship from the Uehara Memorial Foundation.

## Author contributions

A.P.J. and D.A.L. designed the experiments. A.P.J. prepared stimulus set. K.W.K., D.B.T.M., E.N.W., B.E.R. and D.A.L. performed animal surgeries. K.W.K., D.B.T.M. and B.E.R. run fMRI experiments and analysis. K.W.K, A.P.J. and E.N.W. collected electrophysiology data. K.W.K, A.P.J. and D.A.L. performed analysis of neuronal data. K.W.K and D.A.L. wrote the paper. All the authors contributed to editing of the manuscript. D.A.L. supervised the study.

