## Supplemental Figures for "Delayed suppression normalizes face identity responses in the primate brain"

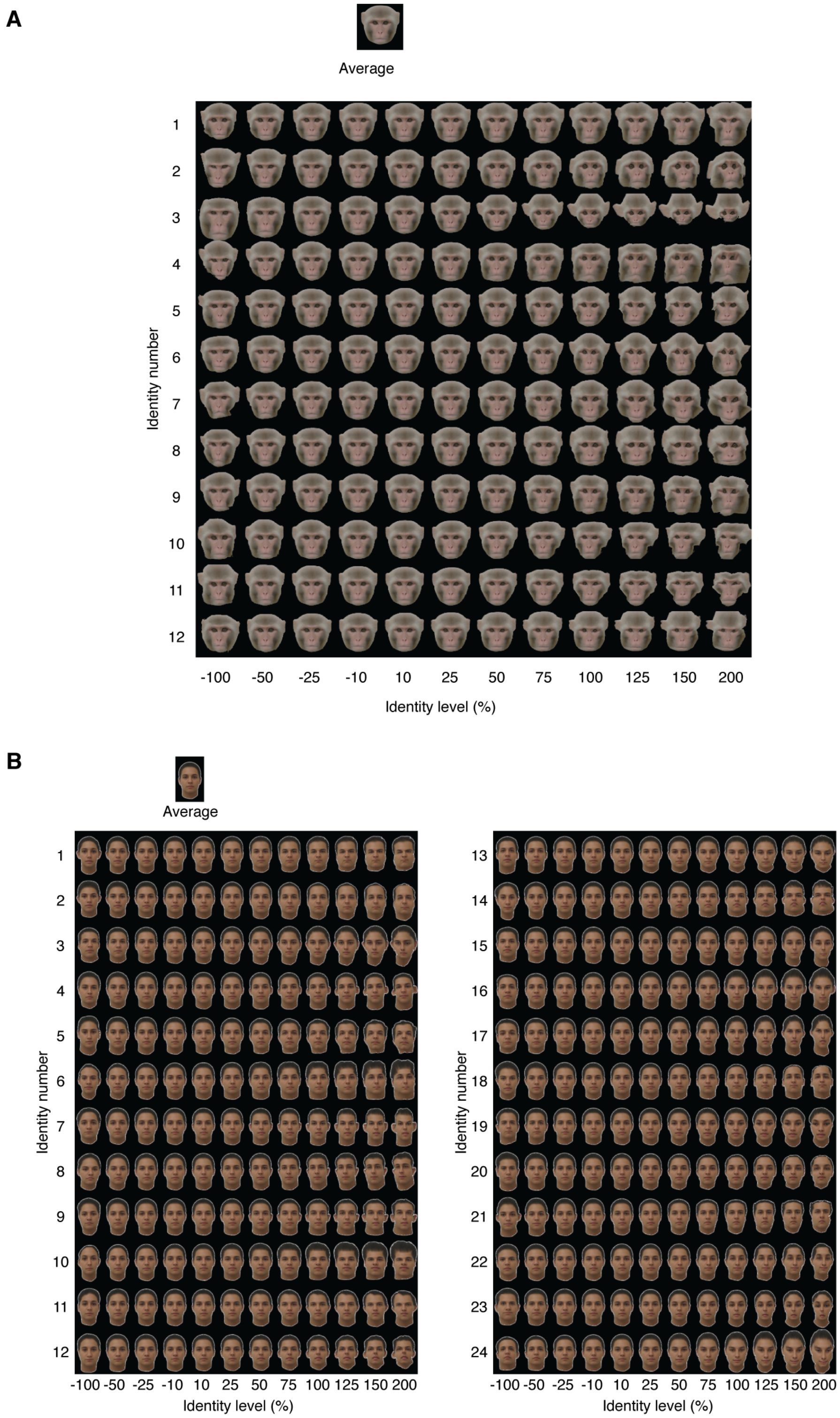


**Figure S1. All face stimuli used for examining neuronal tuning to identity level. (Related to Figure 1).**

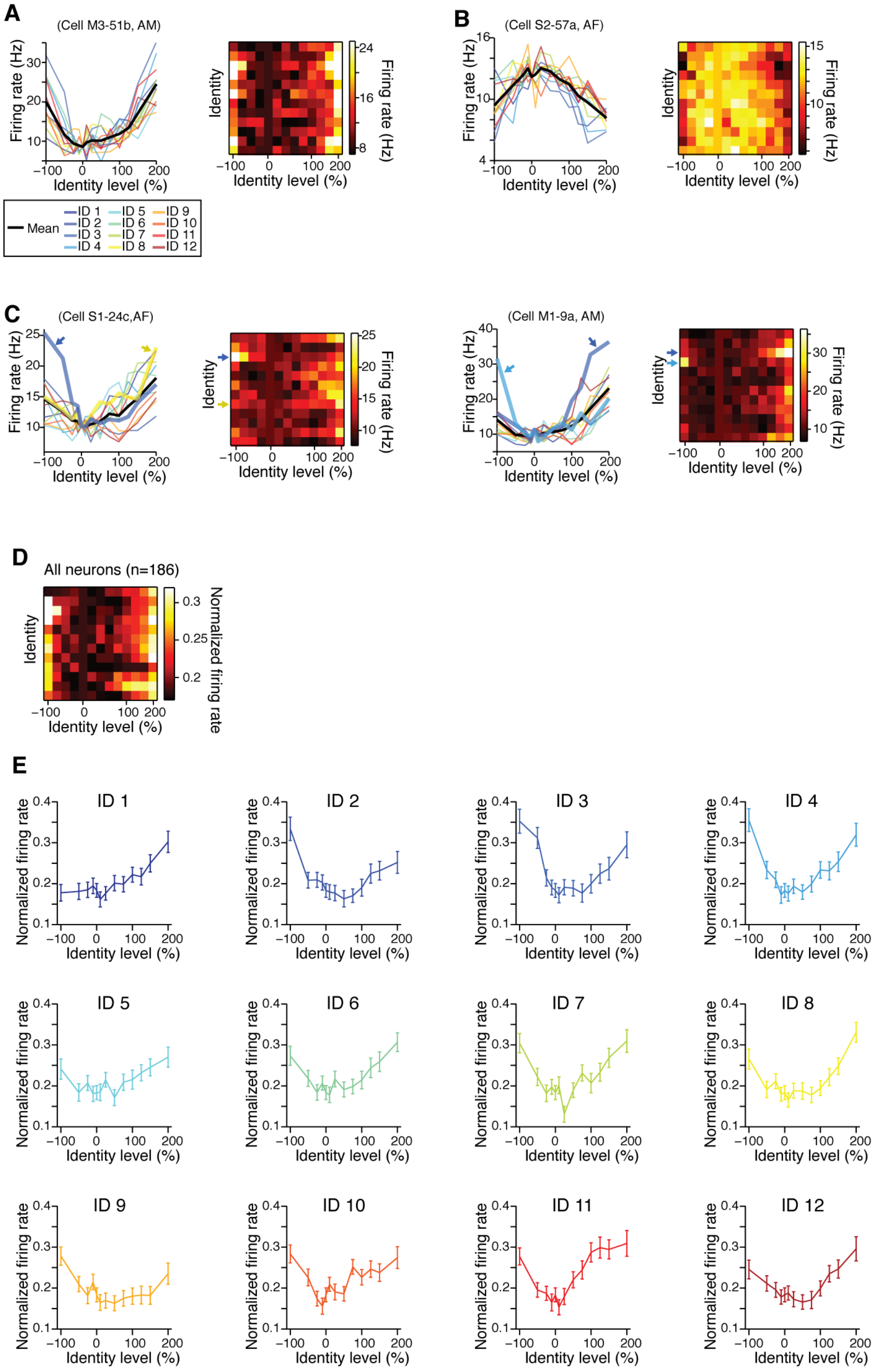


**Figure S2. V-shaped tuning of each identity trajectories (Related to Figure 3).**

(**A**-**C**) Additional example neurons that showed V-shaped tuning. (**A**) A neuron showing V-shaped tuning along most of the identity trajectories. (**B**) A neuron showing Λ-shaped tuning along most of the identity trajectories. (**C**) Neurons showing knee-shaped tuning (arrows) along some identity trajectories. Note that these neurons showed V-shaped tuning when averaged across identities (mean, black line in the left panels). (**D**-**E**) Population-averaged tuning of each identity trajectories. (**D**) A heatmap showing low-firing tendency at the average face (0% identity). (**E**) Tuning to identity level in each identity trajectories. Most of the identity trajectories showed V-shaped tuning pattern, indicating that the tuning shape did not depend on specific identity. Data are represented as mean ± SEM.

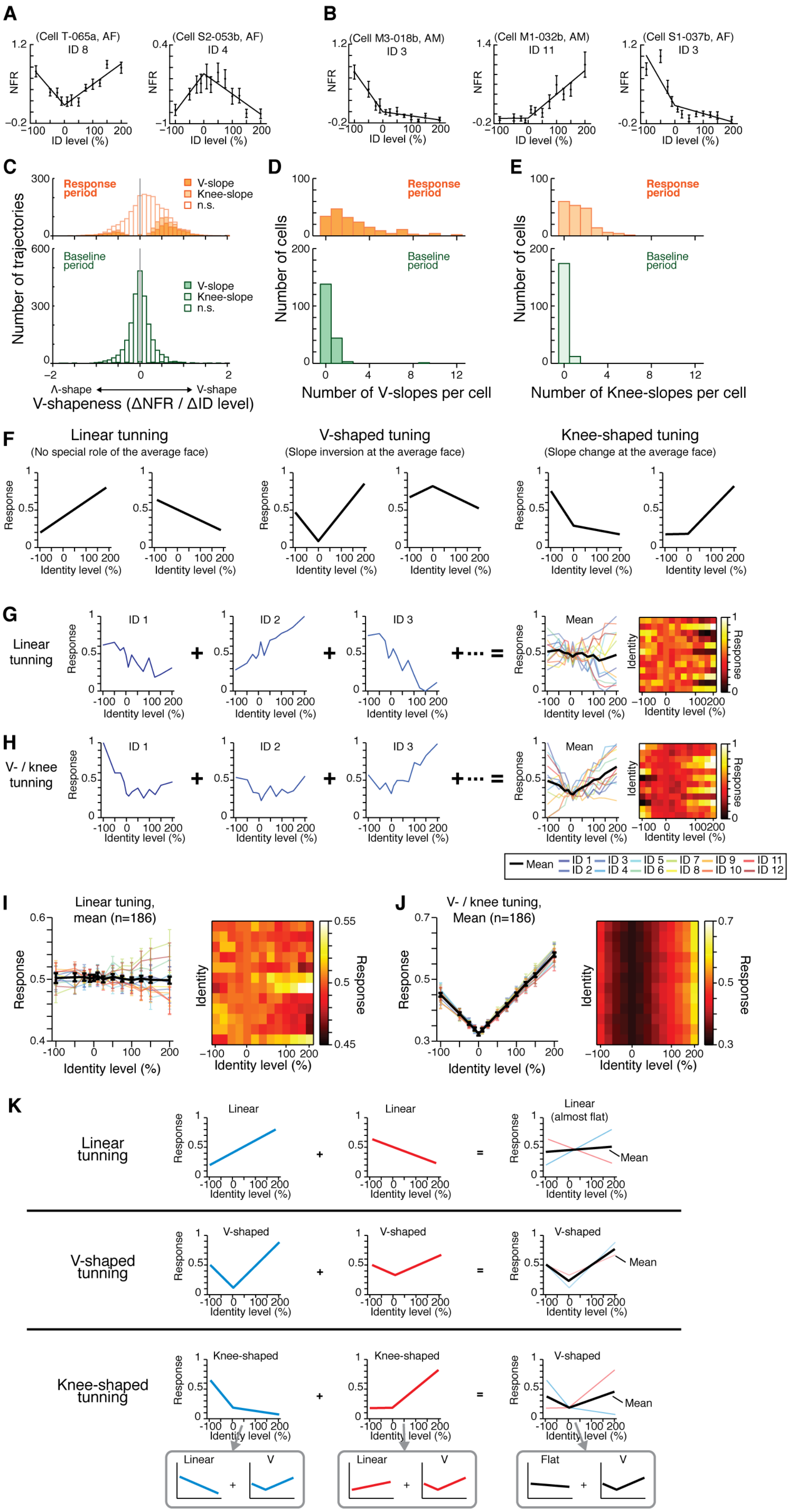
A
(Cell T-065a, AF)
ID 8
(Cell S2-053b, AF)
ID 4
B
(Cell M3-018b, AM)
ID 3
(Cell M1-032b, AM)
ID 11
(Cell S1-037b, AF)
ID 3
NFR
ID level (%)
C
Response period
V-slope
Knee-slope
n.s.
Baseline period
Number of trajectories
Λ-shape
V-shape
V-shapeness (ΔNFR / ΔID level)
D
Number of cells
Number of V-slopes per cell
E
Number of Knee-slopes per cell
F
Linear tunning
(No special role of the average face)
V-shaped tuning
(Slope inversion at the average face)
Knee-shaped tuning
(Slope change at the average face)
Response
Identity level (%)
G
Linear tunning
H
V- / knee tunning
ID 1
ID 2
ID 3
Mean
Identity
Mean
ID 1
ID 2
ID 3
ID 4
ID 5
ID 6
ID 7
ID 8
ID 9
ID 10
ID 11
ID 12
I
Linear tuning, mean (n=186)
J
V- / knee tuning, Mean (n=186)
K
Linear tunning
Linear
Linear (almost flat)
V-shaped tunning
V-shaped
Knee-shaped tunning
Knee-shaped
Linear
V
Flat

**Figure S3. Tuning of each identity trajectories (Related to Figure 3).**

(**A**-**E**) Regression analysis of the tuning shape for each identity trajectories. (**A**) Examples of V-slope identity trajectories with regression lines (V-shaped trajectory on the *left* and Λ-shaped trajectory on the *right*). (**B**) Examples of Knee-slope trajectories with regression lines. (**C**) Distributions of V-shapeness during response (*top*) and baseline (*bottom*) periods. V-shapeness was significantly biased toward positive values during the response period ($p$ < 0.001, Wilcoxon's sign-rank test). V-slope was defined as a slope that was significantly different and had a opposite sign between the caricature (right) and anti-face (left) side of the two regression lines. Knee-slope was defined as a slope that was significantly different but had the same sign between the caricature and anti-face side of the two regression lines. During the response period, 486 (21.8%) trajectories showed v-slope and 241 (10.8%) trajectories showed knee-slope. During the baseline period, most (2161, 96.8%) of trajectories did not show significant difference between regression slopes. (**D**) Number of V-slopes per neuron, which is larger during the response (*top*) than baseline (*bottom*) period ($p$ < 0.001, Wilcoxon's sign-rank test). (**E**) Number of knee-slopes per neurons, which is larger during the response (*top*) than baseline (*bottom*) period ($p$ < 0.001, Wilcoxon's sign-rank test). (**F**-**K**) Simulation examining effect of averaging of knee tuning across identity trajectories on the mean tuning shape. (**F**) Schematic drawings of linear (*left*), V-shaped (*middle*) and knee-shaped (*right*) tuning those were created for simulation in (**G**)-(**J**). (**G**) A simulated responses of a neuron that had linear tuning in each identity trajectories (ID 1, ID 2, ID 3, … on the *left* side), which were created from random slopes and random noises. The averaged response of the linear tuning resulted in linear-shaped tuning (mean, black line on the *right*). (**H**) A simulated responses of a neuron that had v-shaped (e.g. ID 2 on the *left*) or knee-shaped (e.g. ID 1 and ID 3 on the *left*) tuning in each identity trajectories, which were created from random positively-biased right-side slopes, random negatively-biased left-side slopes and random noises. The averaged response of the V/knee tuning resulted in V-shaped tuning (black line, *right*). (**I**) Simulated population-averaged tuning based on the linear tuning, showing flat tuning. (**J**) Simulated population-averaged results based on the v-/knee tuning, showing v-shaped tuning. (**K**) An interpretation of the simulation results. When each identity trajectory shows linear tuning (*upper*), the slopes of the trajectories are canceled out after averaging and result in nearly-flat tuning of mean response. When each identity trajectory shows V-shaped tuning (*middle*), the mean response results in V-shaped tuning. When each identity trajectory shows knee-shaped tuning (*lower*), the mean response results in V-shaped tuning. This is also shown in a different manner at the bottom of (**K**): Since the knee-shaped tuning can be represented as a combination of linear and V-shaped tuning (see also Fig. 7), the components of linear tuning are canceled out after averaging while the components of V-shaped tuning form the mean V-shaped tuning after averaging. Data are represented as mean ± SEM.

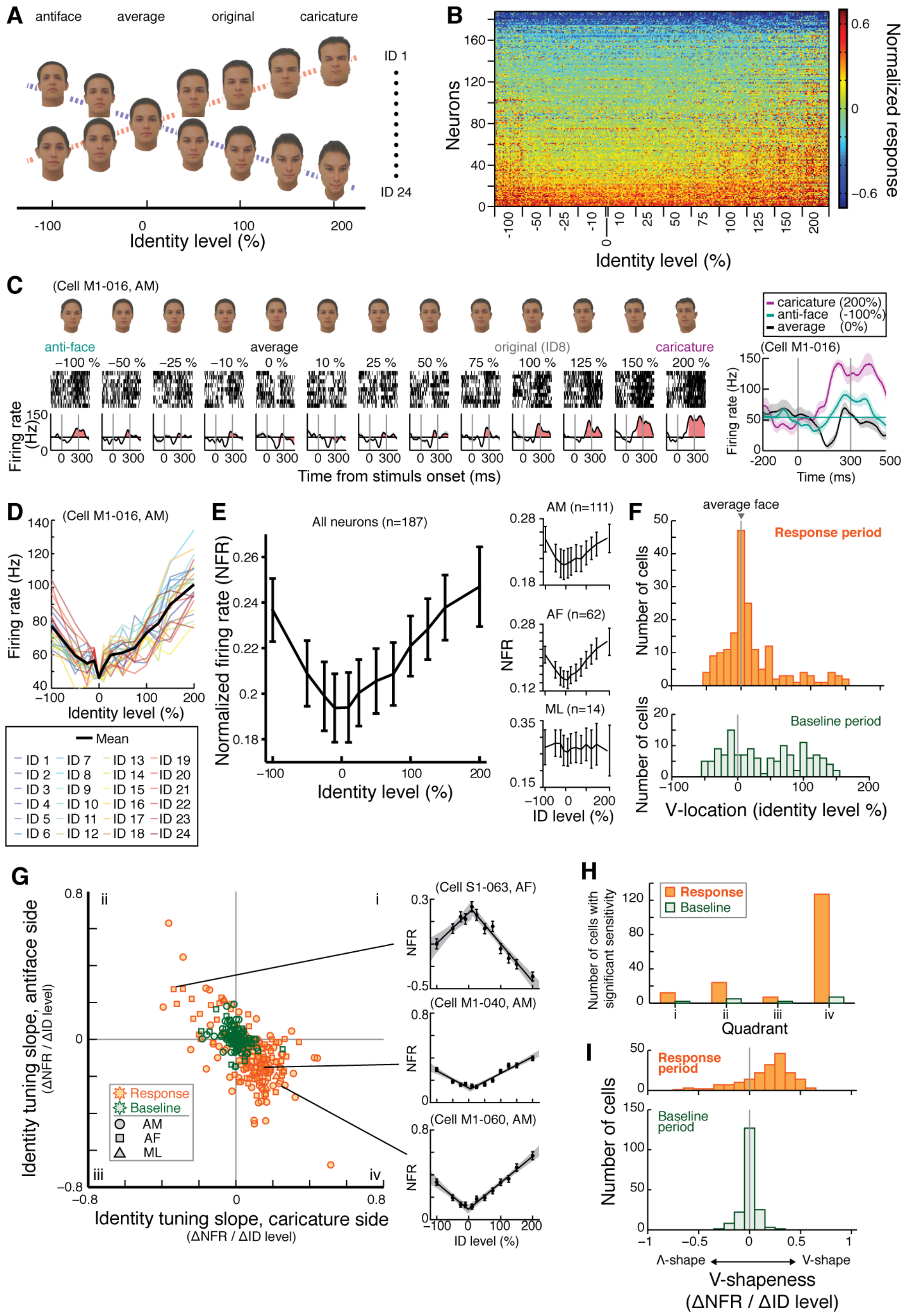

**Figure S4. V-shaped tuning for human faces (Related to Figure 3).**

(**A**) Stimulus set. An average face (0% identity level) was first created by averaging 24 faces. Then faces were morphed between the average and each of the 24 original (100%) identities, creating caricatures (200%) and antifaces (-100%) which have exaggerated and opposite facial features of the original, respectively. (**B**) Responses of face-selective neurons (n=187) for human face stimuli. Neurons tended to respond more to faces with high-identity level. (**C**) Response of a cell in an identity trajectory, exhibiting smallest response to the average face. The response increased in both caricature and antiface side, with distance from the average. (**D**) Response of the neuron in (**C**) for all the identity trajectories. The mean response showed "V-shaped" tuning pattern along the identity level. (**E**) Mean response of all the face-selective neurons, showing characteristic V-shaped tuning. Insets, mean of each of AM, AF and ML patches. (**F**) V-location distributions during response (*top*) and baseline (*bottom*) period. Many neurons had V-location around 0% identity level (average face): during the response period, 110 (58.8%) neurons showed V-location within ±25% identity level, while only 45 (24.1%) did during the baseline period. (**G**) Regression slopes. During response period, slopes in caricature side tended to be anti-correlated with slopes in antiface side, indicating V-shaped or Λ–shaped tuning. (**H**) Number of neurons with significant sensitivity, counted in each quadrant of scatter plot in (**G**). During response period, majority (127 , 67.9%) of neurons were in the fourth quadrant (V-shape) and second highest number (24, 12.8%) of neurons were in the second quadrant (Λ-shape). (**I**) V-shapeness distributions during response (*top*) and baseline (*bottom*) period. V-shapeness was significantly biased toward positive values during the response period ($p$ < 0.001, Wilcoxon's sign-rank test). During the response period, 150 (80.2%) neurons showed significantly different regression slopes, while only 11 (5.9%) did during the baseline period. Data are represented as mean ± SEM.

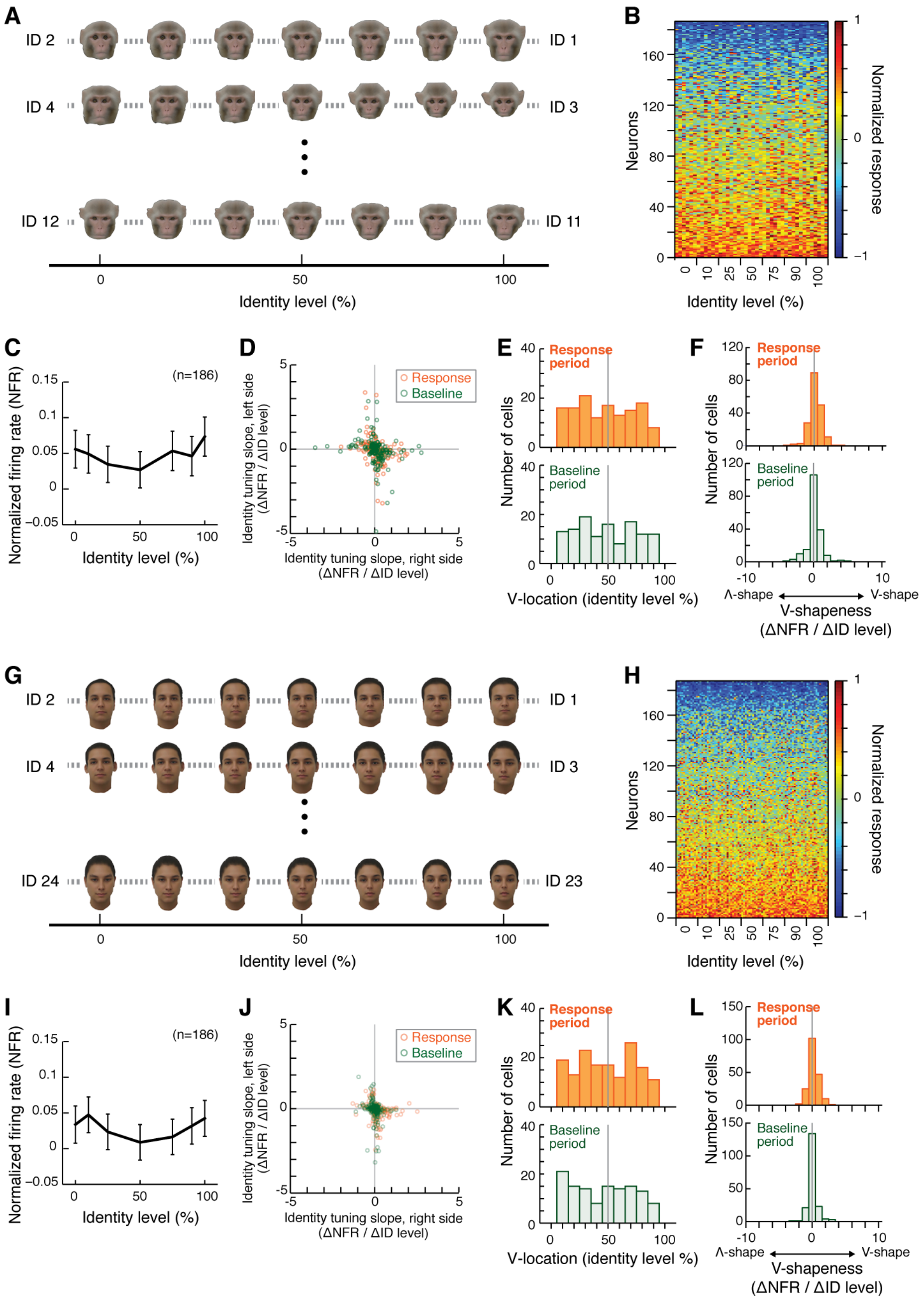

**Figure S5. Morphing between original identities (Related to Figure 3).**

(**A**) Stimulus set. Original 12 faces were divided into 6 pairs and then intermediate morphs were created between the paired original identities. Here in this figure, 0% and 100% were one of the original faces and 10-90% were morphed faces. (**B**) Responses of the face-selective neurons (n=186) for monkey face stimuli. Neurons were the same to that in Fig. 2 and 3. There was no obvious tendency of response for identity level. (**C**) Mean response of all the face-selective neurons, showing no obvious tuning for identity level ($p$ = 0.92, one-way ANOVA). (**D**) regression slopes. Distribution of neurons in the response period overlapped with that in the baseline period. (**E**) V-location distributions during response (*top*) and baseline (*bottom*) period, showing no clear peak. (**F**) V-shapeness distributions during response (*top*) and baseline (*bottom*) period. Only 20 (10.7%) and 11 (5.9%) neurons showed significantly different regression slopes during the response and baseline period, respectively. (**G-L**) Responses of neurons for morphed human faces. Conventions are the same as in A-F. No obvious tuning for identity level ($p$ = 0.93, one-way ANOVA) as shown in (**I**). V-shapeness distributions in (**L**) showed that there were only 34 (18.2%) and 9 (4.8%) neurons that had significantly different regression slopes during the response and baseline period, respectively. Data are represented as mean ± SEM.

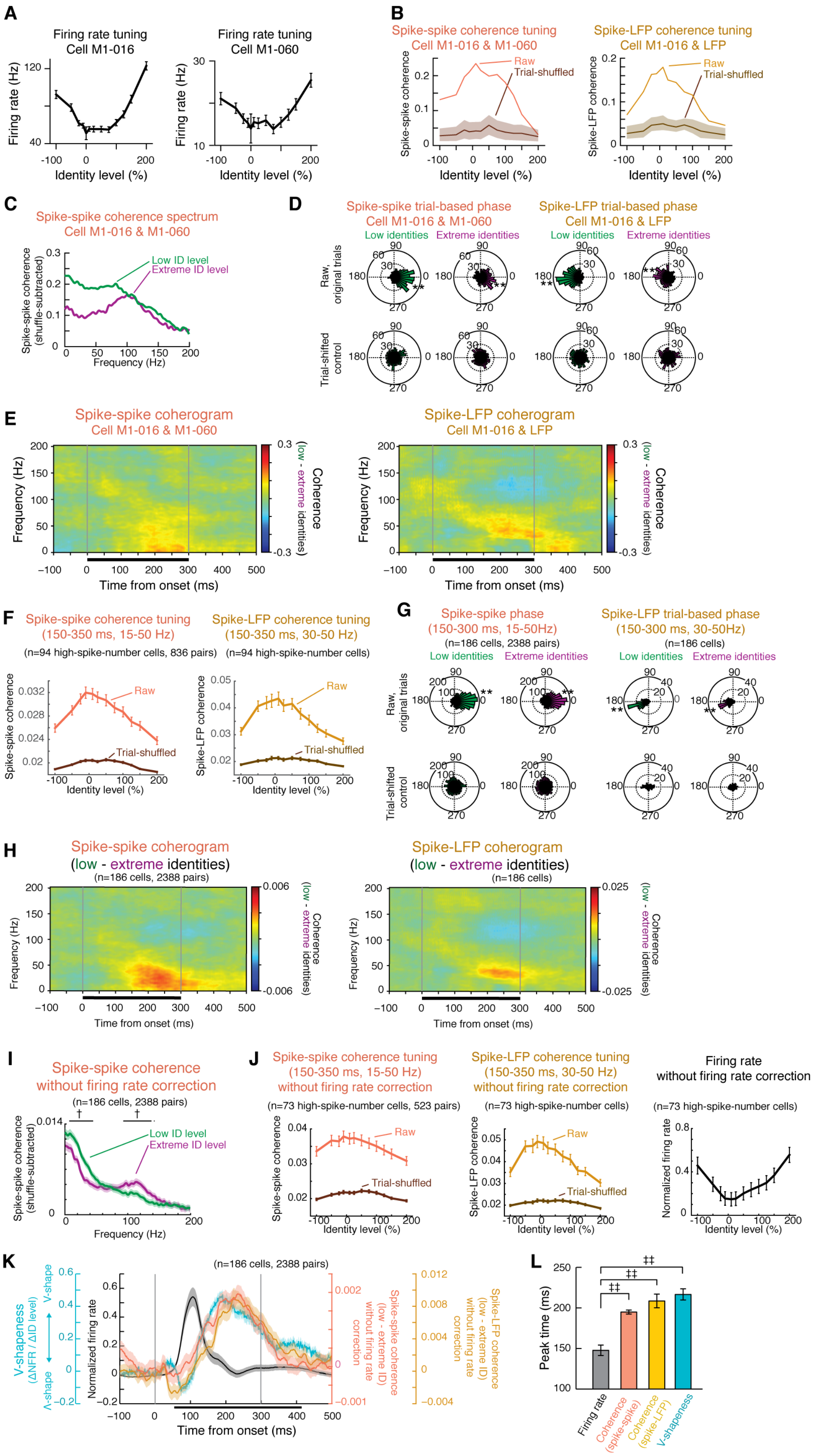
A
Firing rate tuning
Cell M1-016
Firing rate tuning
Cell M1-060
Firing rate (Hz)
Identity level (%)
B
Spike-spike coherence tuning
Cell M1-016 & M1-060
Spike-LFP coherence tuning
Cell M1-016 & LFP
Raw
Trial-shuffled
Spike-spike coherence
Spike-LFP coherence
Identity level (%)
C
Spike-spike coherence spectrum
Cell M1-016 & M1-060
Low ID level
Extreme ID level
Spike-spike coherence (shuffle-subtracted)
Frequency (Hz)
D
Spike-spike trial-based phase
Cell M1-016 & M1-060
Spike-LFP trial-based phase
Cell M1-016 & LFP
Low identities
Extreme identities
Raw, original trials
Trial-shifted control
E
Spike-spike coherogram
Cell M1-016 & M1-060
Spike-LFP coherogram
Cell M1-016 & LFP
Frequency (Hz)
Time from onset (ms)
Coherence (low - extreme identities)
F
Spike-spike coherence tuning
(150-350 ms, 15-50 Hz)
(n=94 high-spike-number cells, 836 pairs)
Spike-LFP coherence tuning
(150-350 ms, 30-50 Hz)
(n=94 high-spike-number cells)
Raw
Trial-shuffled
Spike-spike coherence
Spike-LFP coherence
Identity level (%)
G
Spike-spike phase
(150-300 ms, 15-50Hz)
(n=186 cells, 2388 pairs)
Spike-LFP trial-based phase
(150-300 ms, 30-50Hz)
(n=186 cells)
Low identities
Extreme identities
Raw, original trials
Trial-shifted control
H
Spike-spike coherogram
(low - extreme identities)
(n=186 cells, 2388 pairs)
Spike-LFP coherogram
(low - extreme identities)
(n=186 cells)
Frequency (Hz)
Time from onset (ms)
Coherence (low - extreme identities)
I
Spike-spike coherence
without firing rate correction
(n=186 cells, 2388 pairs)
Low ID level
Extreme ID level
Spike-spike coherence (shuffle-subtracted)
Frequency (Hz)
J
Spike-spike coherence tuning
(150-350 ms, 15-50 Hz)
without firing rate correction
(n=73 high-spike-number cells, 523 pairs)
Spike-LFP coherence tuning
(150-350 ms, 30-50 Hz)
without firing rate correction
(n=73 high-spike-number cells)
Firing rate
without firing rate correction
(n=73 high-spike-number cells)
Raw
Trial-shuffled
Spike-spike coherence
Spike-LFP coherence
Normalized firing rate
Identity level (%)
K
(n=186 cells, 2388 pairs)
V-shapeness (ΔNFR / ΔID level)
V-shape
Λ-shape
Normalized firing rate
Spike-spike coherence (low - extreme ID) without firing rate correction
Spike-LFP coherence (low - extreme ID) without firing rate correction
Time from onset (ms)
L
Peak time (ms)
Firing rate
Coherence (spike-spike)
Coherence (spike-LFP)
V-shapeness

**Figure S6. Coherence showing higher values for the average face (Related to Figure 5).**

(**A**-**E**) Example neuronal pair that showed low-frequency coherence during processing of low identity stimuli (**A**) V-shaped firing rate tuning for identity level of a simultaneously recorded pair of neurons. (**B**) Spike-spike coherence between the neuronal pair (*left*) and spike-LFP coherence of one of the neurons (*right*). Coherences were higher at lower identity levels, shaping inverted v-shaped (Λ-shaped) pattern. Shaded region of trial-shuffled control, 95% confidence interval. (**C**) Frequency spectrum of spike-spike (*left*) and spike-LFP (*right*) coherences. Trial-shuffled control was subtracted from raw coherence. Low identity stimuli tended to elicit higher coherence, especially in lower frequency range. (**D**) Trial-based phase relationship of spike-spike coherence at γ/β range (15-50 Hz, *left*) and spike-LFP coherence at γ range (30-50 Hz, *right*). **, $p < 0.001$, Rayleigh test. (**E**), Spike-spike (*left*) and spike-LFP (*right*) coherogram, showing difference of low identity levels from extreme identity levels. Faces with lower identity levels tended to elicit larger coherence, starting at 150 ms after stimulus onset in lower frequency bands. (**F**-**H**) Coherences during late response period showing higher value for the average face. (**F**) Population-averaged spike-spike (*left*) and spike-LFP (*left*) coherences at each identity level, which were calculated for neurons with enough number of spikes. Coherences tended to be higher at lower identity levels. The dataset of spike-spike coherence was the same to that of Fig. 5D, but raw and trial-shuffled were plotted separately here. (**G**) Distributions of trial-averaged mean spike-spike (*left*) and spike-LFP (*right*) phase of coherences. **, $p < 0.001$, Rayleigh test. (**H**) Population-averaged spike-spike (*left*) and spike-LFP (*right*) coherograms, showing differences between low and extreme identity levels. Cohrence differences showed a peak during later response phase starting at 150 ms after stimulus onset, in γ/β range (15-50 Hz, spike-spike coherogram on the left) and in γ range (30-50 Hz, spike-LFP coherogram on the right). (**I**-**L**) Coherences calculated without spike-drop control, showing essentially similar results to that of Figure 5. (**I**) Frequency spectrum of population-averaged coherences. †, $p<0.05$ (paired *t*-test with Bonferroni's correction). (**J**), Population-averaged spike-spike (*left*) and spike-LFP (*middle*) coherences at each identity level, and firing rate tuning (*right*). Coherences tended to be higher at lower identity levels while firing rate showed opposite pattern. (**K**) Time course of normalized spike firing rate, v-shapeness, spike-spike and spike-LFP coherence calculated with sliding windows. (**L**) Peak time of firing rate, v-shapeness and coherences. ‡‡, $p<0.01$ (paired t-test). Data are represented as mean ± SEM.

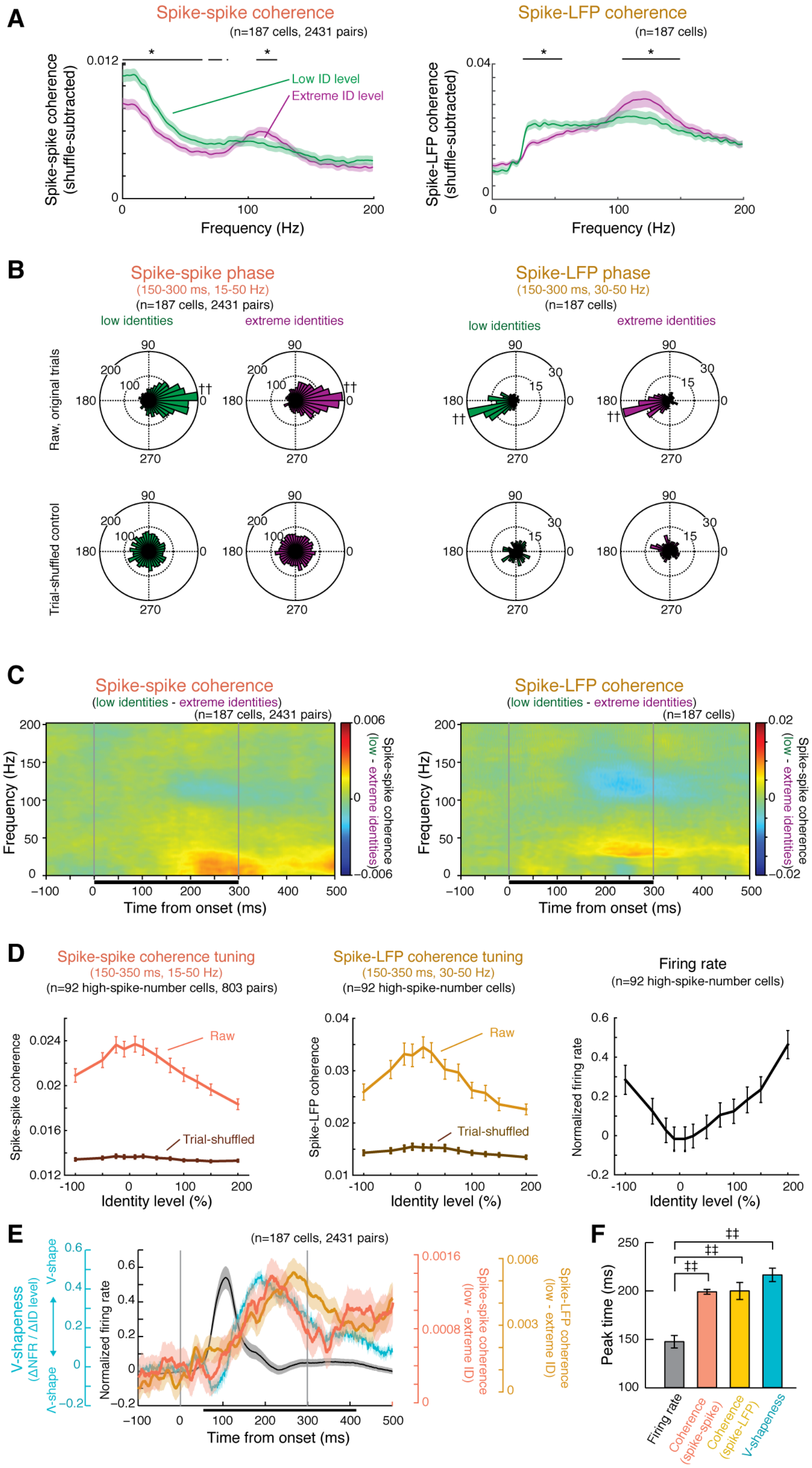


**Figure S7. Coherences for human face stimuli (Related to Figure 5).**

(**A**) Frequency spectrum of population-averaged coherences. *, p<0.05 (paired *t*-test with Bonferroni's correction). (**B**) Distributions of trial-averaged mean phase of coherences. ††, $p < 0.001$, Rayleigh test. (**C**) Population-averaged coherograms showing differences between low and extreme identity levels. (**D**,

Population-averaged spike-spike (*left*) and spike-LFP (*left*) coherences at each identity level, and firing rate tuning. Coherences tended to be higher at lower identity levels while firing rate showed opposite pattern. (**E**) Time course of normalized spike firing rate, v-shapeness, spike-spike and spike-LFP coherence calculated with sliding windows. (**F**) Peak time of firing rate, v-shapeness and coherences. ‡‡, $p<0.01$ (paired t-test). Data are represented as mean ± SEM.
